# Independent origins of fish endothermy converge on a developmental regulatory signature

**DOI:** 10.64898/2026.06.24.734300

**Authors:** Dahiana Arcila, Fernando Melendez-Vazquez, Julian Gallego-Garcia, Emily Ignatoff, Jessica Zhong, Wayne Pfeiffer, Ricardo Betancur-R

**Affiliations:** Scripps Institution of Oceanography, University of California San Diego, La Jolla, CA 92093, USA; American Museum of Natural History, New York, NY 10024, USA; San Diego Supercomputer Center, University of California San Diego, La Jolla, CA 92093, USA

## Abstract

Why independent origins of the same complex physiological trait repeatedly produce similar body forms and physiological changes remains central in evolutionary biology. Endothermy, the internal production and retention of metabolic heat, evolved at least four times in ray-finned fishes, providing natural replicates for convergent genomic signatures. We present a chromosome-scale analysis of three transitions (opah, tunas, and swordfish), including a new chromosome-level genome of the rare, charismatic Pacific oarfish, analyzed with 31 other teleost genomes. The strongest signal is regulatory: of 253,680 conserved noncoding elements, 577 are rate-accelerated in endothermic lineages, with 67 accelerated in all three, exceeding matched ectothermic controls and enriched near developmental transcription factors and Wnt-signaling genes (e.g., *irx1a*, *irx5a*, *her9*, and *lmo1*). These elements overlap zebrafish developmental enhancers more than expected by chance but are not tied to genes emphasized by expression or coding-selection studies of endothermic lineages, marking a regulatory layer distinct from that metabolic layer. This convergence is part of a broader mosaic: endothermic lineages also share transition-biased substitution and convergent duplication signatures, including excess tandem duplications and lineage-specific gene-family expansion, whereas chromosome organization and protein-coding sequence change little. Endothermic convergence therefore leaves its clearest signal in regulatory remodeling, alongside shifts in substitution bias and gene-family evolution.

## Main

Convergent evolution offers a powerful lens for identifying the genomic correlates of complex phenotypes. When similar traits arise independently across distantly related groups, shared molecular signatures can reveal how evolutionary constraints and selection interact to shape biological diversity. Among vertebrates, endothermy (the capacity to generate and retain metabolic heat) represents one of the clearest examples of physiological convergence^1,2^, having evolved independently in mammals, birds, some sharks, and a small number of ray-finned fishes^3,4^. In teleosts, endothermy is restricted to three distantly related clades but represents four independent origins: Lampriformes (opahs), Scombriformes (tunas and butterfly kingfish, two distinct origins), and Carangaria (swordfishes and billfishes)^5,6^. Each independent origin is associated with a distinct suite of anatomical, ecological, and metabolic adaptations^7,8^: red-muscle and visceral endothermy in scombrids, where continuous aerobic swimming generates heat to support high cruising speeds^5^; cranial endothermy via heater organs derived from eye muscles in butterfly kingfish, sailfishes, marlins, and swordfishes^2,5^; and whole-body endothermy, found only in opahs^6,9^. The repeated emergence of this rare and energetically demanding trait provides a natural opportunity to investigate whether similar genomic features recur across independent evolutionary origins^3,4,10–13^.

Chromosome-scale genome assemblies now enable integrated analyses of regulatory evolution, lineage-specific gene-family dynamics, and genome organization, including synteny, for dissecting complex traits and convergent adaptations^14,15^. High-contiguity references have revealed gene-expression modules underpinning limb regeneration in brittle stars^16^, antiviral immune novelties in bats^17^, the genomic history of vertebrate whole-genome duplication using hagfish assemblies^18^, ancient gene linkages anchoring metazoan evolution^19^, and regulatory elements repeatedly accelerated in birds with convergently reduced tarsus length^20^. In ray-finned fishes, chromosome-scale data have revealed genomic changes underlying pelvic-fin loss in percomorphs^21^, toxin evolution in stonefishes^22^, temperature adaptation in snakeheads^23^, and hypoxia resistance in cyprinids^24^. Previous work on fish endothermy has identified candidate genes, physiological pathways, and ecological signatures, including convergent positive selection on *carnmt1* and *dcaf6* across multiple endothermic scenarios in vertebrates^3^ and shared selection on *pkmb* and *ryr1a* between tunas and billfishes^25^. Most studies, however, have focused on individual lineages, limited genomic regions, or transcriptomic data, and have rarely incorporated chromosome-scale architecture or regulatory evolution. The extent to which noncoding regulatory elements, substitution biases, gene-duplication dynamics, and chromosome structure show parallel changes across independent endothermic transitions remains unexplored.

Here, we test whether convergent regulatory evolution is a central genomic signature of repeated endothermic transitions in ray-finned fishes, and how it sits alongside other axes of genomic change. We generated a chromosome-level genome assembly for the Pacific oarfish (*Regalecus russellii*), the closest ectothermic relative of the whole-body endothermic opah (*Lampris incognitus*), and integrated it with 31 teleost genomes spanning whole-body, regional, and cranial endothermy. Using a reference-free whole-genome alignment framework, we first apply a phylogenetically explicit test for convergent rate acceleration in conserved noncoding elements (PhyloAcc-GT), benchmarking these signals against matched ectothermic non-sister trios. We then embed this regulatory analysis in a broader comparative framework that examines chromosome-scale architecture, substitution spectra, gene-family dynamics, gene-duplication architecture, and protein-coding selection across the same three transitions. Rather than seeking a singular genetic “mechanism” of endothermy, we treat convergent regulatory evolution as the anchoring signal and test whether it is accompanied by a consistent mosaic of other recurrent genomic features. Our results support a mosaic model of genomic convergence in which complex phenotypic traits leave more consistent signatures in how genomes evolve than in the specific changes they encode.

### A new oarfish genome anchors the comparative framework

We generated a chromosome-level reference genome for the Pacific oarfish (*Regalecus russellii*), the first such assembly for an ectothermic lampriform and the closest extant ectothermic relative of the whole-body endothermic opah (*Lampris incognitus*; Fig. 1A). The polished assembly spans 746 Mb with 96% Actinopterygii BUSCO completeness, and Hi-C anchoring recovered 24 chromosome-scale scaffolds consistent with the karyotype (2n = 48); assembly statistics, repeat content, and heterozygosity are reported in Methods (Fig. S1; Table S1).

**Fig. 1.**
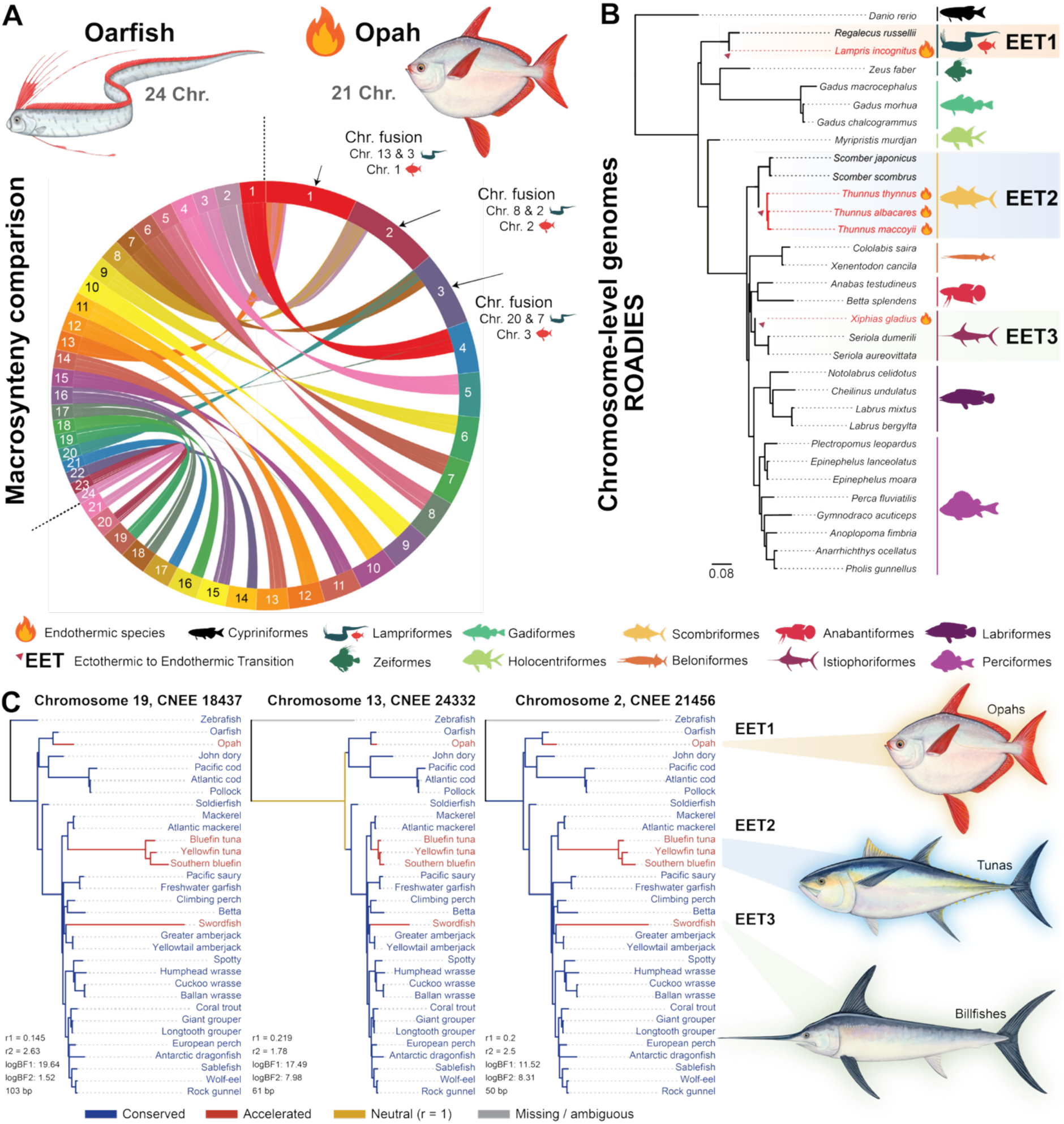
Phylogenomic framework and macrosynteny across independent endothermic transitions. **(A)** Macrosynteny comparison from the hierarchical alignment (HAL) generated with Progressive Cactus between the 24 chromosomes of oarfish (*Regalecus russellii*) and the 21 chromosomes of opah (*Lampris incognitus*); chromosome silhouettes denote fusions in *L. incognitus,* and colors match the corresponding *R. russellii* chromosomes. **(B)** Species tree estimated from 32 chromosome-level genome assemblies using ROADIES (∼32,000 single- and multi-copy gene trees). Trees inferred from UCEs and single-copy nuclear exons under IQ-TREE are provided in Fig. S2 and Dataset S1. Transitions from ectothermy to endothermy are marked based on ancestral state reconstruction (Fig. S3): EET1, opah lineage; EET2, tunas; EET3, swordfish. Endothermic species are marked with a flame symbol; each order is denoted by a unique color and a representative species silhouette. **(C)** Representative examples of convergently accelerated conserved noncoding elements (EARs) at three loci (Chr 19, CNEE 0497; Chr 11, CNEE 24332; Chr 2, CNEE 21416). For each element, branches of the species tree are colored by inferred substitution-rate state from the Bayesian acceleration model: conserved (blue), accelerated (red), neutral (r = 1; gold), or missing/ambiguous (gray), showing independent rate acceleration on the three endothermic lineages (opah, EET1; tunas, EET2; billfishes, EET3). Per-element Bayes factors (logBF1, logBF2) and element lengths (bp) are indicated.

We reconstructed species relationships from three independent genomic datasets (reference-free genome-wide single- and multi-copy gene trees via ROADIES^26^, ultraconserved elements^27^, and single-copy nuclear exons), which were topologically congruent for our focal taxa (Fig. 1B; Fig. S2; Dataset S1). To identify independent origins of endothermy, ancestral-state reconstruction on a time-calibrated tree of 1,051 ray-finned fishes^3^ strongly supported ectothermy at all nodes subtending the three focal transitions (nodal probability ≥ 0.996; Fig. S3), recovering three independent transitions: whole-body endothermy in opah (EET1), regional red-muscle endothermy in tunas (EET2), and cranial endothermy in billfishes (EET3). The cranially endothermic butterfly kingfish, a putative second transition in Scombridae, is not sampled here.

### Convergent acceleration of conserved noncoding elements accompanies independent endothermic transitions

Because many complex phenotypes evolve through changes in gene regulation rather than protein sequence alone^28–29^, we tested whether the three independent origins of endothermy are associated with recurrent shifts in regulatory constraint. We analyzed conserved nonexonic elements (CNEEs) using PhyloAcc-GT, a Bayesian method that detects lineage-specific changes in substitution rate while accounting for gene-tree discordance^30^. From a Progressive Cactus^31^ multiple-genome alignment, we identified 253,680 CNEEs (>20 bp), of which 577 qualified as endothermic accelerated regulatory regions (EARs) in at least one endothermic lineage. EARs were defined by support for target-lineage acceleration over both the conserved null and unconstrained acceleration models (logBF1 >10 and logBF2 >1; Fig. S20). A stricter logBF2 >3 cutoff reduced the set to 534 EARs, and logBF2 >5 to 477, without changing the convergence pattern (Fig. S7).

Per-lineage EAR counts were 237 in opah, 379 in swordfish, and 193 in tunas (Fig. 2A), partitioned into lineage-specific (132 in opah, 228 in swordfish, 52 in tunas) and shared accelerations. Pairwise overlaps were 14 (opah and tunas), 60 (tunas and swordfish), and 24 (opah and swordfish). A total of 67 EARs were accelerated in all three lineages (Fig. 2B,C). The observed convergence greatly exceeded null expectations: permuting the endothermic focal set against 10,000 random ectothermic non-sister trios matched for total target-branch length and taxonomic composition yielded a null median of 0 triple-convergent EARs (empirical P < 10⁻⁴; Fig. 2B) and 0 doubly-convergent EARs (P < 10⁻⁴).

**Fig. 2.**
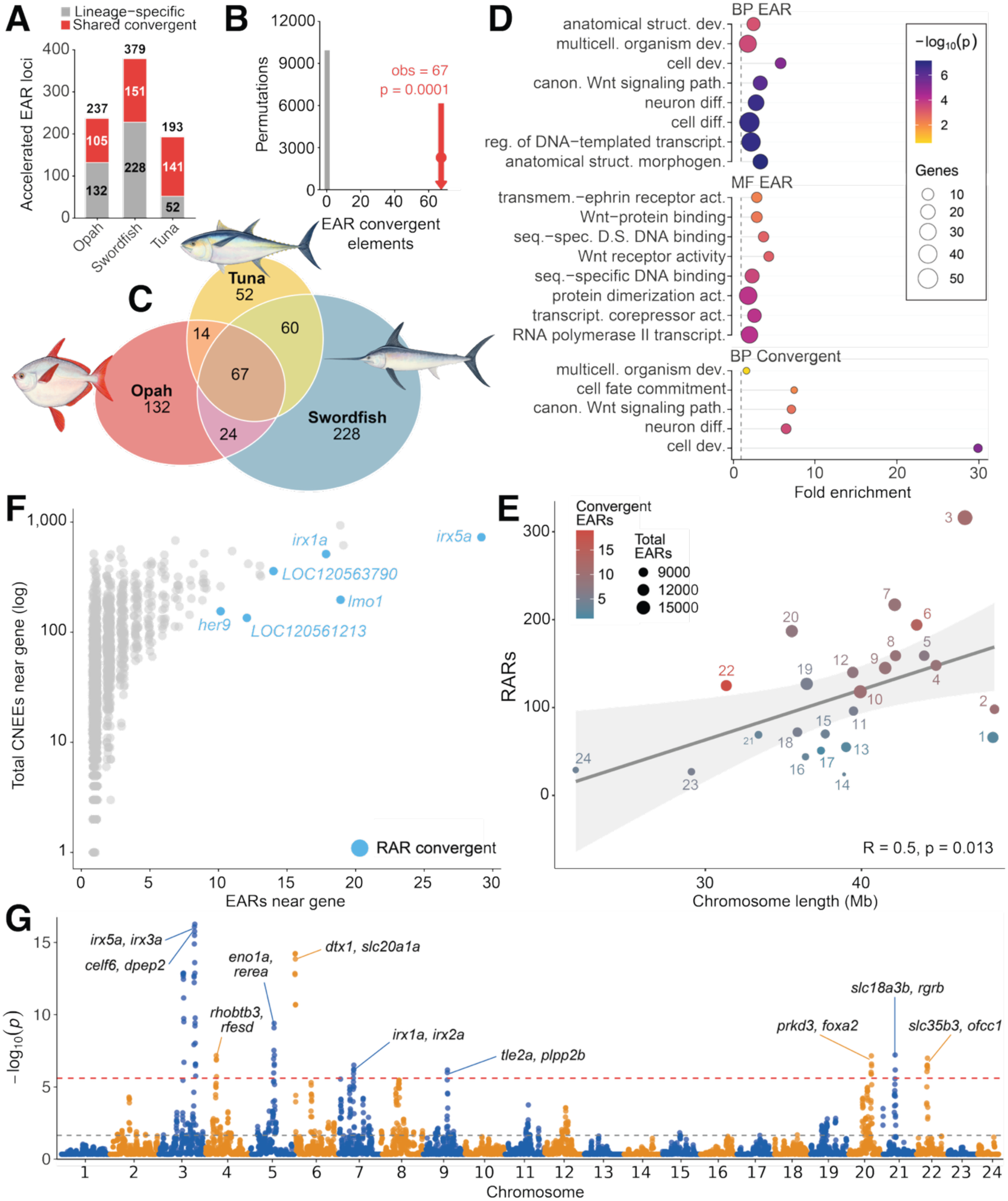
Convergent acceleration of conserved noncoding elements across independent origins of endothermy. **(A)** Counts of rate-accelerated regulatory regions (EARs; logBF1 >10 and logBF2 >1) per endothermic lineage, partitioned into lineage-specific (gray) and shared-convergent (red): opah (EET1), 237; swordfish (EET3), 379; tunas (EET2), 193. **(B)** Permutation test for triple convergence. The observed number of EARs accelerated in all three endothermic lineages (n = 67) exceeds the null distribution of 10,000 random ectothermic non-sister trios matched for branch length, taxonomic composition, and CNEE density (gray; median null = 0; empirical P <10⁻⁴). **(C)** EAR overlap among the three endothermic lineages. Triple-convergent set, n = 67; pairwise overlaps: opah and tunas = 14, tunas and swordfish = 60, opah and swordfish = 24; lineage-specific counts: opah = 132, swordfish = 228, tunas = 52 (recovering the per-lineage EAR totals in panel A). **(D)** Functional enrichment for the full EAR set and the triple-convergent subset. The convergent subset is dominated by cell development (GO:0048468; ∼30-fold enrichment), neuron differentiation, canonical Wnt signaling, cell-fate commitment, and multicellular organism development. **(E)** Per-gene aggregation of accelerated CNEEs. Each point is one annotated gene; x-axis, number of accelerated CNEEs in cis; y-axis, total CNEE density in cis (log₁₀). Loci passing FDR ≤0.05 with ≥3 accelerated EARs and ≥2 convergent EARs in cis are labelled. **(F)** Per-chromosome relationship between scaffold length (Mb) and target-accelerated CNEE count (Pearson R = 0.5, P = 0.013). **(G)** Genome-wide Manhattan plot of per-CNEE convergence P-values, with chromosomes alternating in color. The red dashed line marks the genome-wide significance threshold; the gray dashed line marks P = 0.05. The strongest peaks fall on *Perca fluviatilis* chromosomes 3, 5, 6, and 21.

PhyloAcc detects shifts in evolutionary constraint, not positive selection per se: acceleration may reflect relaxed constraint, directional selection, or processes such as GC-biased gene conversion³². Even so, the recurrence of this regulatory signal across three independently derived endothermic lineages, and its excess over matched ectothermic trios (permutation-based P <10⁻⁴), suggests that similar selective regimes or cellular environments have repeatedly reshaped developmental regulatory landscapes.

### Convergent EARs show enhancer-associated signatures at developmental loci

Accelerated CNEEs were unevenly distributed, with several 1-Mb windows on *Perca fluviatilis* chromosomes 3, 5, 6, and 21 showing strong local enrichment (P < 10⁻¹⁵; Fig. 2G). Convergent elements also lay at slightly greater median distances from transcription start sites than lineage-specific EARs (log10 distance ∼4.5 vs. 4.2; Fig. S8), consistent with distal regulatory activity. Gene-level enrichment, controlling for local CNEE density, recovered six loci with at least three accelerated CNEEs at FDR ≤ 0.05, led by the iroquois homeobox genes *irx5a* (29 accelerated CNEEs) and *irx1a* (20), together with *lmo1*, *sall1a*, *her9*, and *ofcc1* (Fig. 2E; Figs. S16, S17, S19; these loci and their flanking accelerated CNEEs are shown in Fig. 4D). Gene Ontology enrichment of the genes nearest convergent EARs, relative to the whole genome, was dominated by cell development (GO:0048468; fold enrichment ∼30), neuron differentiation, canonical Wnt signaling, and cell-fate commitment, with molecular-function terms for sequence-specific DNA binding and transcription-factor activity (Fig. 2D; Table S10; Fig. S18). About 45% of the 577 EARs lay within 10 kb of an annotated gene and 81% within 50 kb (median distance ∼12 kb), consistent with cis-regulatory effects on nearby developmental loci.

Because CNEEs as a class are themselves clustered and already flank developmental regulators, neither this spatial clustering nor the genome-wide GO enrichment shows that the accelerated subset is functional. We therefore compared EARs to background CNEEs in four location-based tests (FDR corrected) plus a separate variant-effect test. The clearest independent signal was enhancer association: projected onto the zebrafish genome through the whole genome alignment, EARs overlapped developmental enhancer histone marks (H3K27ac and H3K4me1; DANIO-CODE,^72^) more often than a length-matched CNEE background (25.5% vs. 18.6%; permutation P = 0.010, q = 0.042 after correction; Table S21). Enhancer-overlapping EARs again flanked the same developmental regulators, including *irx5a*, *her9*, and the Wnt effector *lef1*. Because these marks derive from zebrafish rather than from the endothermic fishes or their heat-producing tissues, the overlap supports a regulatory role but does not demonstrate activity in opah, tunas, or swordfish.

By contrast, EARs were not preferentially near genes differentially expressed in endothermic muscle (tuna and opah transcriptomic and proteomic datasets^73,74^), near positively selected genes from a prior analysis of ray-finned fish endothermy^3^, or enriched for transcription-factor motifs once sequence length was controlled (all P >0.08; Table S21). Supervised variant-effect prediction with AlphaGenome^75^ found no concentration of large-effect substitutions on endothermic versus ectothermic-sister branches (one-sided P ≈0.09–0.11), consistent with acceleration being distributed across many small-effect changes rather than concentrated in a few high-impact variants. Because AlphaGenome is trained on human and mouse data, these teleost predictions are out-of-distribution and should be interpreted cautiously. In summary, convergent EARs appear to be enhancer-associated elements at developmental loci, a regulatory layer distinct from the metabolic-effector genes identified by prior expression and coding-selection studies.

### Chromosome-scale organization is conserved while substitution spectra shift consistently

We examined large-scale genome organization across the three transitions using Progressive Cactus alignments and halSynteny^33^. Chromosome-scale gene order was broadly conserved between ectothermic-endothermic sister pairs, with most chromosomes retaining extensive collinearity. In the lampriform comparison (EET1), the opah genome contains fewer chromosomes than oarfish (21 vs. 24), reflecting lineage-specific fusions; large syntenic blocks remained intact, indicating that restructuring primarily involved the joining of pre-existing chromosomal segments instead of widespread rearrangement (Fig. 1A). Comparisons within EET2 (tunas vs. mackerels) and EET3 (billfishes vs. jacks) showed even greater structural conservation, with most differences confined to local inversions and short-range translocations (Fig. S4). Large-scale chromosomal rearrangement was therefore not a shared feature of the three transitions.

To assess whether endothermic and ectothermic lineages share genome-wide patterns of sequence evolution, we summarized mutation spectra as log-transformed transition-to-transversion ratios (log(ts/tv)) extracted with halSummarizeMutations and compared them via phylogenetic ANOVA. In the pooled tip-level comparison, endothermic lineages showed higher log(ts/tv) than ectothermic relatives on average (mean Δlog(ts/tv) = 0.18, 95% CI = 0.03–0.33; P = 0.027; Fig. 3A), consistent with a group-level shift toward more transition-biased substitution spectra. However, this pattern was not uniform across the three independent transitions: the contrast was positive in EET1 (opah vs. oarfish) and EET2 (tunas vs. mackerels), but negative in EET3 (swordfish vs. jacks; Table S3). This suggests a pooled endothermy-associated shift in substitution spectrum, not a uniform signal across all three origins. Transition bias reflects long-term substitutional patterns rather than direct estimates of germline mutation rate, and could arise through base composition, context-dependent mutagenesis (for example, CpG deamination), recombination-associated GC-biased gene conversion, or differences in DNA-repair efficiency^28 NONSENSICAL CITATION^; our analyses do not distinguish among these alternatives. Even so, the positive contrasts in two of the three transitions suggest that substitutional bias remains a plausible genomic correlate of endothermy that warrants broader taxonomic sampling.

**Fig. 3.**
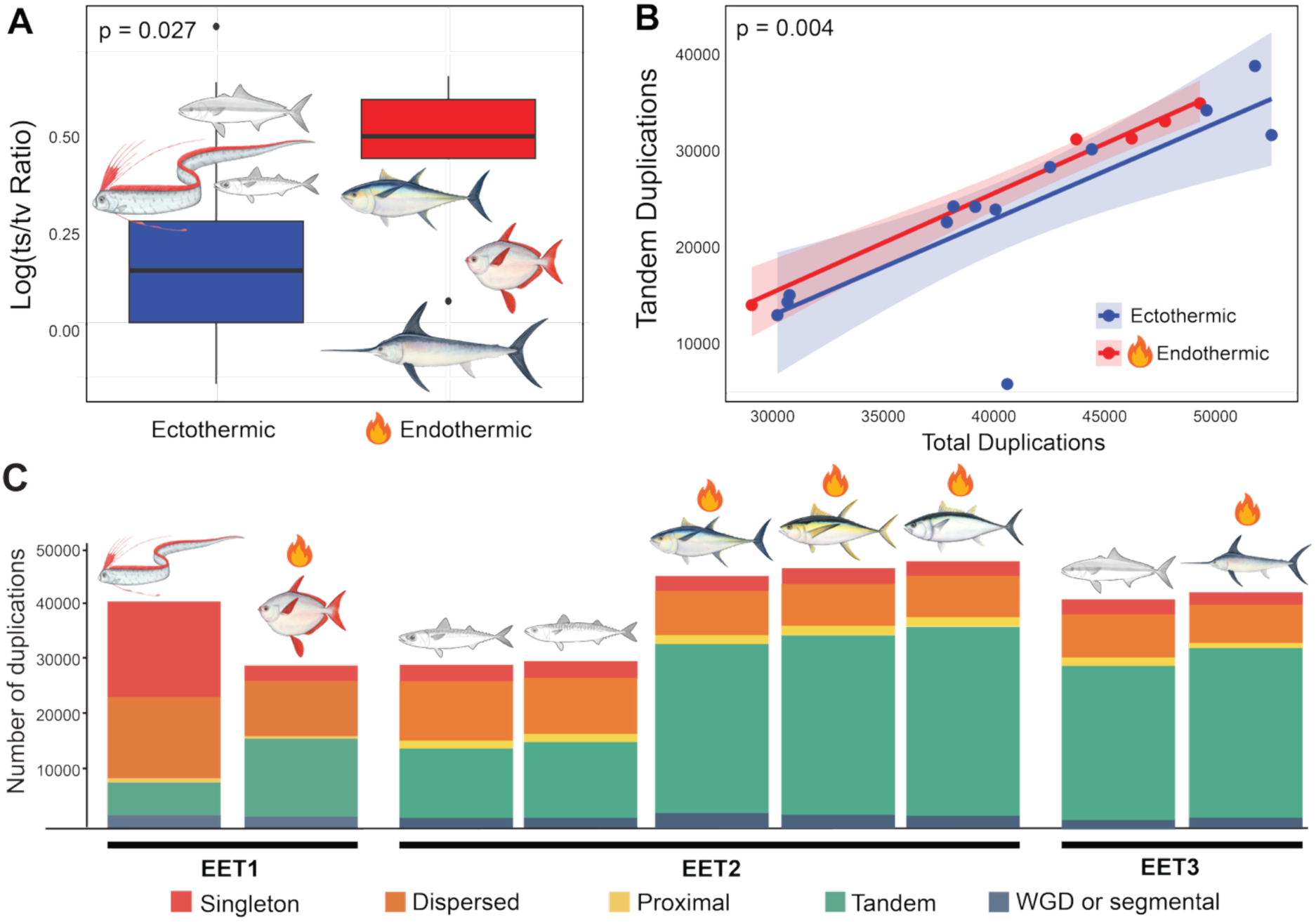
Convergent shifts in substitution spectra and duplication architecture across independent endothermic transitions. **(A)** Phylogenetic ANOVA: endothermic lineages show higher log(ts/tv) than ectothermic relatives (mean Δlog(ts/tv) = 0.18, 95% CI = 0.03–0.33; P = 0.027), consistent with a transition-biased substitution spectrum. **(B)** Phylogenetic generalized least-squares regression of tandem duplications on total duplications in the broader taxon dataset. The tandem-total relationship differed between endothermic (red) and ectothermic (blue) lineages (interaction model vs. additive model, P = 0.004); endotherms showed a shallower fitted slope than ectotherms (interaction beta = −0.80, P = 0.012). **(C)** Per-species duplicate counts by class (singleton, dispersed, proximal, tandem, WGD/segmental), grouped by transition (EET1, opah; EET2, tunas; EET3, swordfish). Flame symbols denote endothermic species.

### Distinctive duplication architecture accompanies convergent gene-family expansion

Because gene duplication is a major source of evolutionary novelty^34^, we asked whether duplication architecture differs between endothermic and ectothermic lineages. Using MCScanX^35^, we classified duplicate genes as tandem, proximal, dispersed, or whole-genome/segmental (Table S4). Across all three transitions, endothermic lineages showed elevated proportions of tandem duplications relative to their ectothermic counterparts, respectively: *L. incognitus* (48.2%) vs. *R. russellii* (14.5%); *Thunnus* species (mean ∼69%) vs. *Scomber* species (mean ∼45%); *X. gladius* (71.3%) vs. *S. aureovittata* (66.5%, a smaller difference in the same direction). To test whether this pattern extended beyond the three focal contrasts, we fit phylogenetic generalized least-squares models of tandem duplication count as a function of total duplications (Fig. 3B). Tandem counts increased with total duplications, but the relationship differed between thermal groups: an interaction model fit better than an additive model (likelihood-ratio test, P =0.004), and the negative interaction term (β =-0.80, P =0.012) indicated a shallower increase in endothermic lineages. At the mean total-duplication level of the sampled taxa, however, endotherms still had higher fitted tandem counts than ectotherms. This pattern suggests different scaling of tandem duplications with overall duplication burden, rather than a steeper endothermic increase or a uniformly higher tandem-duplication share across the broader phylogenetic sample. The repeated elevation of tandem-duplicate fractions across the three focal contrasts indicates that tandem duplication may be one component of genomic differentiation associated with endothermic lineages.

We then analyzed gene-family expansions and contractions on the full phylogeny with CAFE5^36^ on 22,665 gene families. Endothermic lineages showed higher numbers of significantly expanded gene families than their ectothermic counterparts (per-family CAFE p <0.05; Fig. 4A). GO enrichment of expanded families recovered convergent enrichment for muscle structure and contraction, calcium homeostasis, oxidative and steroid metabolism, and cellular stress response across the three transitions (Fig. S22B; Tables S6–S11; Figs. S5–S6, S9–S12). Despite these shared functional themes, the specific gene families contributing to enrichment differed substantially among lineages, and no gene family was significantly expanded across all three transitions under a strict transition-level criterion. A relaxed, tip-level screen recovered 24 orthogroups with significant positive expansion in opah, swordfish, and at least one tuna lineage (Table S14). This overlap did not exceed chance: in a size-matched comparison against non-sister ectothermic triplets, the number of families jointly expanded across the three endothermic lineages fell at the median of the ectothermic distribution (≈ 1.0-fold, empirical P = 0.50), indicating no excess convergence in gene-family expansion beyond lineage-specific change. At a coarse functional level, these 24 families span several themes plausibly relevant to endothermy, including cytoskeletal and extracellular-matrix remodeling (myocardin-related transcription factor B, plexin-A2-like), neural and cell-communication functions (neuroligin 4 X-linked a, transmembrane protein 106B-like), transcriptional and signaling regulation (tyrosine-protein kinase Fer, PDE9A), and metabolic or homeostatic processes (FXYD6-like) such as ion balance and heme biosynthesis (Table S14); we treat these categories as hypothesis-generating annotations from gene predictions rather than as formal functional enrichments. Together, these results indicate that gene-family expansion in endothermic lineages proceeds along multiple, largely independent genomic routes rather than through a shared set of convergently expanded families.

**Fig. 4.**
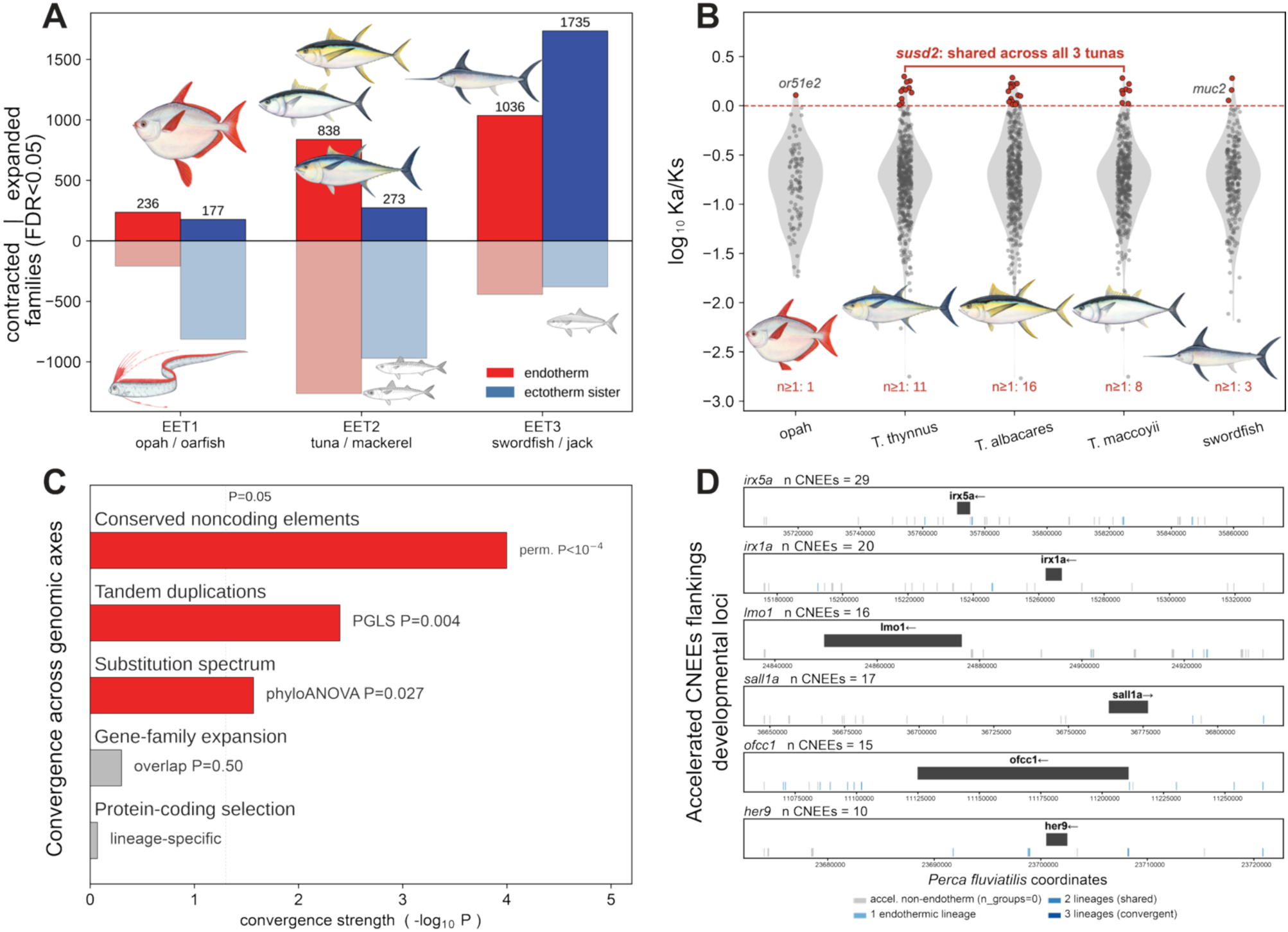
Gene-family and protein-coding evolution are extensive but lineage-specific. **(A)** Gene-family turnover across the three transitions. Bars show the number of significantly expanded (upward) and contracted (downward) gene families (CAFE5, per-family P < 0.05) for each endothermic lineage (red) and its ectothermic sister (blue): EET1 (opah, *Lampris incognitus*, versus oarfish, *Regalecus russellii*), EET2 (tuna, *Thunnus thynnus*, *T. albacares*, *T. maccoyii*, versus mackerel, *Scomber japonicus*), and EET3 (swordfish, *Xiphias gladius*, versus jack, *Seriola aureovittata*). **(B)** Distributions of Ka/Ks for collinear gene pairs (MCScanX) in the endothermic-lineage genomes (opah, three *Thunnus* species, swordfish). Each point is one collinear pair (red, Ka/Ks ≥ 1; grey, Ka/Ks < 1), and the number of candidate pairs per lineage is small (opah 1, *Thunnus* 8–16, swordfish 3). No candidate reached Ka/Ks ≥ 1 in all three independent transitions. *susd2*, a regulator of cell proliferation, was the single candidate shared across the three tunas, whereas *or51e*2 (opah) and *muc2* (swordfish) were lineage-specific. **(C)** Convergence scorecard summarizing the strength of convergence (−log₁₀ P) across the genomic axes examined here: conserved noncoding elements (rate-accelerated regulatory regions; permutation P < 10⁻⁴), tandem-duplication architecture (PGLS P = 0.004), substitution spectrum (phylogenetic ANOVA P = 0.027), gene-family expansion (triplet-overlap permutation P = 0.50), and protein-coding selection (lineage-specific). The dotted line marks P = 0.05; bars to its left indicate no detectable convergence. Values summarize tests reported separately in the text and are placed on a common axis for visual comparison only, not as a single combined test. **(D)** In-silico validation of convergent EARs as developmental enhancers. Accelerated conserved noncoding elements (CNEEs) flanking six marquee developmental loci (*irx5a*, *irx1a*, *lmo1*, *sall1a*, *ofcc1*, *her9*) in *Perca fluviatilis* coordinates; each tick is a CNEE, with grey marking elements accelerated only on non-endothermic lineages and blue shades marking acceleration in one, two, or three endothermic lineages. These loci accumulate many accelerated CNEEs, most on non-endothermic branches, illustrating the regulatory landscape around the convergent developmental regulators. Quantitative results of all validation tests, including the enhancer-mark overlap, are given in Fig. S21, Table S21. Per-lineage expansion-and-contraction counts on the phylogeny and the gene-level functional enrichment of expanded families are shown in Fig. S22.

### Protein-coding convergence is sparse, episodic, and lineage-specific

To test whether independent origins of endothermy share protein-coding changes, we examined both genome-wide patterns of selection in conserved genomic regions and a targeted set of candidate genes nominated by prior work. We screened collinear gene pairs for elevated nonsynonymous-to-synonymous substitution-rate ratios (Ka/Ks) using MCScanX. The analysis identified 118,666 collinear genes out of 205,166 total annotated protein-coding genes (57.84%), defining a deeply conserved syntenic framework in which genes are expected to experience strong functional constraint. Despite this large conserved set, very few collinear genes showed Ka/Ks ≥ 1: a single gene in opah (or51e2; 0.0008%), 8–16 genes per *Thunnus* species (including shared elevation on *susd2*, a regulator of cell proliferation, and lineage-specific signals on *dcaf7*), and three in *X. gladius* (including *muc2*, involved in epithelial protection; Tables S15–S19; Fig. 4B). Across all transitions, candidate loci with Ka/Ks ≥1 represented fewer than 0.02% of the conserved gene set. Whereas convergence was strong for conserved noncoding elements throughout the genome, it was not detectable for gene-family expansion or protein-coding selection (Fig. 4C). Targeted MEME re-tests of two prior candidates (*carnmt1* and *dcaf6*^3^) confirmed episodic positive selection at a small number of codon sites in EET2 and EET3 (Supplementary Methods; Figs. S14–S15; Tables S12, S13). Together with the Ka/Ks scan, these results indicate that protein-coding adaptation in endothermic lineages, where present, is localized and lineage-specific rather than recurrent across transitions.

## Discussion

Three of the four independent origins of endothermy in ray-finned fishes share a common regulatory footprint^3^. Across opah, tunas, and swordfish, genome-wide scans identify recurrent acceleration of conserved noncoding elements, including a substantial set shared across independent endothermic lineages and exceeding matched ectothermic expectations. This regulatory convergence sits within a broader mosaic of endothermy-associated genomic change: shifts in substitution spectra, distinctive duplication architectures, heterogeneous gene-family expansions sharing broad functional themes, and sparse, episodic coding adaptation. Rather than identifying a single molecular “mechanism” of endothermy, our results suggest that repeated evolution of this complex trait leaves its most consistent signature in regulatory remodeling, with other genomic axes acting as complementary, partly independent components of convergence.

The convergent EARs point specifically to developmental gene-regulatory networks. Their strongest gene-level enrichments occur at developmental transcription factors and canonical Wnt-signaling components, including *irx1a*, *irx5a*, *her9*, *lmo1*, *sall1a*, and *ofcc1*, implicating repeated remodeling of body-plan and cell-fate regulatory networks as a central axis of endothermic evolution. GO enrichment of the nearest genes is dominated by cell development, neuron differentiation, canonical Wnt signaling, and cell-fate commitment. Beyond accelerated sequence evolution alone, these elements show enhancer-associated support: projected onto the zebrafish genome, they overlap developmental enhancer histone marks more than length-matched background^72^, and enhancer-overlapping EARs flank the same developmental regulators, including *irx5a*, *her9*, and the Wnt effector *lef1*. This signal is consistent with the hypothesis that the morphological and physiological innovations underpinning endothermy (red-muscle repositioning, countercurrent vascular networks, enlarged aerobic fields, and warmed retinal and neural tissue for sharper vision at depth^2,5,9,13^) arose through repeated tinkering with conserved developmental regulatory networks rather than through parallel recruitment of a narrow stress-response or protein-homeostasis module. A genome-wide rerun across all three transitions also revised the inference of an earlier single-chromosome pilot, which had nominated stress-response pathways; this shift from pilot to genome-wide analysis strengthens the developmental interpretation reported here.

The other axes are best read as complementary signals. Chromosome-scale architecture is conserved, with no large-scale rearrangement shared across transitions, so genome restructuring is not a convergent feature of endothermy. By contrast, substitution spectra and duplication architecture point to finer-scale genomic shifts that may share an upstream context: higher metabolic rates and warmer tissues, with attendant oxidative stress (though acting through different mechanisms). Warmer tissues can increase spontaneous cytosine deamination, producing transition mutations and fitting the transition-biased spectrum we observe, although life-history traits, population size, or recombination could also contribute³²,⁷⁷. Elevated tandem-duplication fractions instead suggest repeated local changes in gene copying, favoring fine-scale gene-family remodeling rather than convergence on the same duplicated genes³⁴,⁷⁸. One possibility is that oxidative and replication stress associated with active metabolism makes template switching more likely, increasing the formation of tandem copies. Finally, protein-coding adaptation is sparse, lineage-specific, and concentrated at a small number of previously implicated candidate loci such as *carnmt1* and *dcaf6*³,²⁵.

A further reason these axes may resist convergence is the nature of the phenotype itself. Endothermy is not one character but a set of alternative solutions, from whole-body heat retention in opah to cranial heating of the eyes and brain in billfishes and regional red-muscle heating in tunas. When functionally equivalent outcomes can be assembled from different anatomical and physiological parts, the mapping from genotype to phenotype is many-to-one, and independent origins need not recruit the same genes even as they converge on the same function^76^. This view predicts that the clearest convergent signal should reside in the shared developmental-regulatory scaffolding common to these solutions, whereas the gene-family and protein-coding changes that implement each solution should remain largely lineage-specific, as we observe. Collectively, these axes outline a mosaic model of genomic convergence in which the regulatory signal anchors a set of complementary, partly independent changes.

A few caveats remain. Substitution spectra reflect long-term evolutionary patterns and are not a direct readout of specific mutational processes without context-dependent analyses. Duplication calls also depend on assembly and annotation quality. Regulatory acceleration nominates candidate elements rather than demonstrating their function: cross-species chromatin data place them at developmental enhancers, but direct tissue-specific expression and chromatin-accessibility assays in the endothermic species will be needed to establish how they contribute to physiological phenotypes. Finally, additional chromosome-scale genomes from unsampled or underrepresented endothermic lineages, including butterfly kingfish²,³, will refine our understanding of how universal these genomic signatures are. In conclusion, by integrating chromosome-scale structure, substitution bias, duplication architecture, coding adaptation, and regulatory evolution, we provide a template for studying convergence of key innovations across the tree of life, in which similar selective regimes repeatedly reshape genomes along predictable axes.

## Methods

### Specimen acquisition and ethics

The *Regalecus russellii* specimen used for this study was a beach-stranded individual recovered after washing up in La Jolla, San Diego, California, in August 2024. The specimen was deceased on recovery and required no live-animal handling; no IACUC approval was therefore required. The carcass was stored at –80 °C to limit DNA degradation. A necropsy was conducted shortly after the collection event, during which multiple tissue types and blood samples were obtained. Tissues were flash-frozen on collection and stored at −80 °C until DNA extraction. The remainder of the specimen was formalin-fixed, preserved in isopropanol, and deposited in the Scripps Institution of Oceanography Marine Vertebrate Collection (accession SIO123).

### DNA extraction, genome sequencing, and Hi-C library preparation

High-molecular-weight (HMW) DNA was extracted from flash-frozen muscle and fin tissue at the Sanford Consortium Genomics Core Facility (gCore) using the Nanobind Tissue Big DNA Kit (PacBio) following the manufacturer’s protocol for nucleated vertebrate tissues. Because recovered HMW DNA yields were below library-preparation minima (a consequence of post-mortem degradation in the stranded specimen), genomic DNA was amplified using the REPLI-g kit (Qiagen) prior to SMRTbell library construction with the SMRTbell Prep Kit 3.0 (PacBio). To mitigate the known risks of MDA-associated chimeras, coverage bias, and inflated repeat content, we (i) filtered HiFi reads with the PacBio chimera-detection pipeline (lima + ccs --min-passes 3), (ii) quantified MDA-induced k-mer skew with smudgeplot and KAT, and (iii) verified coverage uniformity across chromosome-scale scaffolds (median coverage ratio across chromosomes within 0.95–1.05; Fig. S1; Table S1). DNA was sheared with Covaris g-TUBEs (2,750 rpm) and HiFi sequencing was performed on the PacBio Revio platform at 300 pM loading concentration. For chromosome-scale scaffolding, Hi-C libraries were prepared using the Arima High-Coverage Animal Tissue protocol and sequenced on an Illumina NovaSeq X Plus system using a single lane of a 25B PE150 flow cell.

### Genome assembly and quality control

Genome assembly followed the protocol established by the Vertebrate Genomes Project within the Genome 10K Consortium (G10K)³⁸. Adapter sequences were removed using Cutadapt v5.0³⁹. K-mer-based analyses estimated genome-wide metrics with Meryl v1.4.1⁴⁰ and GenomeScope2 v2.0.1⁴¹, including genome size, repeat content, heterozygosity, and k-mer-fit graphical summaries (Fig. S1). Primary and alternate assemblies were built with hifiasm v0.25.0 (Hi-C phased mode)⁴²⁻⁴⁴, incorporating both PacBio HiFi and Hi-C reads. Initial assembly quality and completeness were assessed with Merqury v1.3⁴⁰. Haplotype-resolved primary contigs were purged of duplicated segments with purge_dups v1.2.6⁴⁵ and decontaminated against a curated database with Kraken2 v2.1.2⁴⁶. The haplotype-resolved assembly was scaffolded with YaHS v1.2.2⁴⁷ using Arima’s Hi-C mapping pipeline. Scaffolding results were visualised with the CurationPretext pipeline⁴⁸. Assembly metrics were computed with gfastats v1.3.10⁴⁹; BUSCO v5.5.0 against Actinopterygii_odb10⁵⁰ was used to assess gene-content completeness (Table S1).

### Repeat masking and annotation

Transposable elements were identified using Dfam TE Tools v1.93⁵¹ (BuildDatabase + RepeatModeler with the --LTRStruct option) and masked with RepeatMasker (–e rmblast –x small) against the resulting custom library. Genome annotation was performed with BRAKER3 v3.0.3⁵² using protein evidence from the Vertebrata OrthoDB partition and RNA-seq read mappings from publicly available transcriptomes (NCBI SRA accessions to be added at submission).

### Multidataset phylogenomic inference

For phylogenetic reconstruction we expanded sampling to include 31 chromosome-level genomes from NCBI (Table S2) spanning 11 orders of ray-finned fishes. We used three sources of molecular data, chromosome-level genomes, ultraconserved elements (UCEs), and exon markers, to assess topological robustness across marker types. ROADIES²⁶ was run on raw genome assemblies with the flags ‘--mode accurate’ and ‘--converge’ to infer a discordance-aware species tree from ∼32,000 sampled gene trees without requiring annotations or reference orthology. UCEs were extracted following PHYLUCE v1.7.3²⁷ using the Acanthomorph bait set (Acanthomorphs-UCE-1Kv1.fasta); 1,095 UCE loci with 500 bp flanks were aligned with MAFFT v7.471⁵³ and analyzed in IQ-TREE v2.0⁵⁴ under the mixture (MIX) model. Exon mining used nhmmer (HMMER v3.1⁵⁵) against a curated single-copy nuclear locus probe set; 1,105 loci were assembled with Trinity v2.8.5⁵⁶, aligned with MACSE v2.05⁵⁷, and analyzed in IQ-TREE v2.0 with PartitionFinder⁵⁸ and ModelFinder⁵⁹. The previously reported “1,038 single-copy nuclear loci” refers to an earlier mining step; the final tree-building set used 1,105 loci after quality-filtering.

### Ancestral reconstruction of endothermy

Because the 32-taxon tree is biased toward endothermic species (endothermy occurs in ∼40 of ∼36,000 ray-finned fish species (∼0.1%)), we used the calibrated and averaged phylogeny of 1,051 species from Melendez-Vazquez et al.³ to perform ancestral state reconstructions and model fitting. Thermal physiology was coded as a binary discrete trait across the 32 taxa. We compared symmetric and asymmetric transition-rate models using fitDiscrete in geiger v2.0.11⁶⁰ and calculated AIC weights. Based on the best-supported model, we used make.simmap in phytools v2.5-2 to perform stochastic character mapping with 1,000 simulations. Model comparison favored an equal-rates over an all-rates-different model (ER AICw = 0.73 vs. ARD AICw = 0.27), and likelihood-based ancestral-state reconstruction under symmetric rates strongly supported ectothermy at all nodes subtending the three focal transitions (nodal probability ≥ 0.996; Fig. S3), recovering three independent endothermic transitions (whole-body in opah, EET1; regional red-muscle in tunas, EET2; cranial in billfishes, EET3).

### Reference-free genome alignment

Reference-free multiple-genome alignment across the study group was generated with Progressive Cactus v2.9.3³¹. The species tree inferred from ROADIES was used as the guide and soft-masked genome assemblies as input. Cactus was run with default alignment parameters under the GPU-enabled configuration, and the resulting alignment was stored in hierarchical alignment (HAL) format³³ as the common substrate for all downstream comparative analyses, including pairwise and multi-species macrosynteny (halSynteny), mutation-spectrum extraction (halSummarizeMutations), four-fold-degenerate-site sampling (hal4dExtract), and conserved-element detection. Alignment completeness was assessed from per-genome coverage within the HAL.

### Pairwise and multi-species macrosynteny

Quantification of pairwise macrosynteny between ectothermic and endothermic taxa was constructed using halSynteny³³ to extract collinear blocks from the HAL alignment produced by Progressive Cactus. Species-specific karyotype files and link files representing syntenic regions were generated with custom scripts (Datasets S2–S3). Final visualizations were rendered with the Circa synteny visualization software (omgenomics.com/circa/). Additional ribbon-plot visualizations of macrosynteny across EET2 and EET3 species were generated with Oxford Dot Plot (ref. 23). For odp, reciprocal all-versus-all BLAST results identified orthologs across *S. scombrus*, *X. gladius*, *T. thynnus*, and *T. maccoyii*. A backbone of endothermic-specific linkage groups was characterised using strict reciprocal best hits (RBH), then expanded to *S. japonicus*, *T. albacares*, and *S. aureovittata* by converting the backbone to HMMs to recover orthologs in cases of imperfect matches or missing data. A 1% dominance filter (retaining the 99% most significant translocations) was applied to suppress minor noise. Colinear-gene blocks across multiple species (EET2 and EET3) were also identified with MCScanX³⁵; conservation of ≥ 5 consecutive genes was required for block detection. Multi-species visualizations used SynVisio⁶¹. Master collinearity and GFF files are provided in Dataset S4.

### Mutation statistics

We used halSummarizeMutations³³ to extract base- and structure-level differences from the Progressive Cactus alignment for each species pair. Substitutions, transitions, transversions, matches, insertions, deletions, inversions, duplications, and transpositions are tabulated in Table S3. For comparative analyses, we summarized the substitution spectrum as log₁₀(ts/tv) per species and tested for endothermy effects under a phylogenetic ANOVA using the pruned ROADIES tree as the covariance structure (effective n = 3 transition pairs). Effect sizes are reported as mean Δlog(ts/tv) (endotherm − ectotherm) with 95% CIs.

### Identification and classification of gene duplications

Gene duplications were identified with duplicate_gene_classifier in MCScanX³⁵, classifying duplicates as tandem, proximal, dispersed, segmental, or singleton (Table S4) based on sequence similarity (internal BLASTP) and positional information. To test whether the relationship between tandem and total duplications differed between endothermic and ectothermic lineages, we fit phylogenetic generalized linear models (PGLMs) using the pruned species tree as the phylogenetic covariance matrix. For each genome, the proportion of tandem duplications (logit-transformed) was the response variable, total duplication count was a covariate, and thermal strategy (endothermic vs. ectothermic) was included as an interaction term. Significance of model terms was assessed by likelihood-ratio tests.

### Gene-family expansion and contraction

Significantly expanded and contracted gene families were identified across the ray-finned fish phylogeny with CAFE5 v1.1³⁶ on 22,665 gene families. The species tree used for CAFE was derived from our ROADIES phylogeny (Fig. 1B) and time-calibrated by congruification⁶² against the 1,051-species chronogram of Melendez-Vazquez et al.³ using shared taxa as anchors. Orthogroups and single-copy orthologous amino-acid sequences were identified with OrthoFinder v2.5.4⁶³. To reduce numerical instability from extreme family-size differentials, we excluded the 120 gene families (∼0.5%) with the largest copy-number differentials; sensitivity at exclusion thresholds of 60 and 240 yielded qualitatively identical patterns of expansion/contraction (Table S5). CAFE was run under the two-category GAMMA model to estimate λ (turnover) and α (rate-class shape), detecting lineage-specific shifts in family size at p < 0.05. To test whether endothermic lineages shared more expanded families than expected, we compared their size-normalized triple overlap (opah, one *Thunnus* species, swordfish) with all non-sister ectothermic triplets drawn from the three clades. The endothermic overlap fell at the ectothermic median (empirical P = 0.50), indicating no excess convergence in shared gene-family expansion.

### Gene Ontology enrichment

Gene Ontology (GO) enrichment of significantly expanded gene families was performed with clusterProfiler v4.2.2⁶⁴ against species-specific GO mappings annotated by eggNOG-mapper v2.1.12⁶⁵ using the Biological Process domain. Annotations were collapsed to the orthogroup level and supplied to the enricher function (one-sided hypergeometric test with Benjamini–Hochberg correction). Foreground sets were the significantly expanded families per ectothermic–endothermic pair (per-family p < 0.05); the background was all families detected across the dataset. GO term descriptions were extracted with the GO.db Bioconductor package v3.7.7⁶⁶ and visualised as dot plots of the top 30 terms per analysis; redundancy was reduced with REViGO⁶⁷ via rrvgo v1.21.1 using the REViGO dispensability score. The ten most relevant GO terms per species were curated by adjusted p-value, endothermic relevance, and enrichment score.

### Evolutionary rates of collinear genes and site-based selection

Nonsynonymous (Ka) and synonymous (Ks) substitution rates were estimated for each endothermic genome using add_Ka_and_Ks_to_collinearity_Yn00.pl from MCScanX³⁵, with self-BLASTP and MCScanX outputs (see “Identification and classification of gene duplications”) as input. Coding sequences were extracted in FASTA format. Elevated rate shifts were defined at Ka/Ks > 1. Targeted MEME re-tests of two prior candidates (carnmt1 and dcaf6) were conducted as a confirmation of ref. 3; full methods, parameters, and rationale are provided in Supplementary Methods.

### Detection of accelerated conserved noncoding elements and in-silico validation

Conserved nonexonic elements (CNEEs) exhibiting lineage-specific acceleration in endothermic lineages were identified using PhyloAcc-GT³⁰ through a Snakemake-based pipeline (Datasets S6–S7). Input consisted of the HAL alignment of 32 ray-finned fish species generated with Progressive Cactus (see “Reference-free genome alignment”). We scaled the analysis to all 24 autosomes of the *Perca fluviatilis* reference (NC_053111.1–NC_053134.1; an initial pilot restricted to NC_052572.1 was folded into the genome-wide run and is reported in the Supplementary Materials for comparison). Four-fold degenerate sites were extracted with hal4dExtract (HAL v2.3³³), and a neutral substitution model was estimated with phyloFit (PHAST v1.8⁶⁹). Conserved elements were predicted with phastCons and filtered to 20–5,000 bp. Per-element FASTA files were generated, concatenated, and linked to a BED partition file. Elements present in fewer than 15% of taxa were excluded. PhyloAcc-GT was run in batch mode with endotherms as the foreground group, the GT model, species-specific gene trees from IQ-TREE⁵⁴, and ASTRAL⁷⁰ summary trees.

Elements were classified as endothermic accelerated regions (EARs) in a given endothermic lineage when logBF1 (target vs. null) > 10 and logBF2 (target vs. unconstrained) > 1; we also report results at a stricter logBF2 > 3 cutoff. To assess convergence, the observed number of EARs shared by at least two or by all three endothermic lineages was compared to permutation null distributions generated by reassigning the foreground to 10,000 ectothermic non-sister trios drawn from the same alignment and matched for total target-branch length, taxonomic composition, and CNEE density. Empirical P-values were computed as the proportion of permutations reaching or exceeding the observed overlap. Gene-level enrichment was assessed with a Poisson model controlling for local CNEE density; GO-term enrichment of the genes nearest convergent EARs was computed relative to the whole genome. Multiple testing was controlled at FDR ≤ 0.05 (results at FDR ≤ 0.10 are also reported for comparison with prior PhyloAcc literature⁷¹). For visualization, accelerated CNEEs flanking the top gene-level candidate loci were plotted along the *Perca fluviatilis* chromosomes by acceleration tier (number of endothermic lineages accelerated; Fig. 4D).

We then tested the 577 EARs (and the 67 convergent subset) for regulatory function with four location-based analyses, corrected for multiple testing together (Benjamini-Hochberg), plus a separate variant-effect test. Each location-based test compared EARs to background CNEEs by permutation (10,000 resamples): we counted how often EARs fell near, or overlapped, the genes or features of interest using the nearest *Perca fluviatilis* (GCF_010015445.1) gene by midpoint distance, and the P value is the fraction of random background sets matching or exceeding the observed count. (1) Expression: genes differentially expressed in endothermic fishes (tuna muscle and heart RNA-seq, ^73^; opah heat-muscle proteomics, ^74^) were matched to *Perca* by name and tested for proximity to EARs, by nearest-gene membership and by the expression change of nearby genes. (2) Positive selection: genes under positive selection in a prior endothermy study ^3^ were tested for the broad (>=1 scenario) and convergent (>=2) sets. (3) Developmental regulators: a GO-defined developmental and Wnt-signalling gene set (QuickGO, zebrafish) was tested for proximity to EARs. (4) Enhancer overlap: EARs were mapped to zebrafish (danRer11) through the HAL alignment and intersected with DANIO-CODE H3K27ac and H3K4me1 marks^72^, using a length-matched permutation confirmed by Fisher exact test. Two further tests were evaluated outside this corrected family. EAR sequences were scanned for 18 JASPAR vertebrate transcription-factor motifs and compared to a length-matched background (EARs are ∼2× longer than the CNEE pool). Separately, AlphaGenome supervised in-silico mutagenesis^75^ scored the 1,211 convergent-EAR substitutions (ancestral vs. derived allele on endothermic vs. matched ectothermic-sister branches); as a human and mouse-trained model, its teleost predictions are out-of-distribution. Full details in Data Availability.

## Acknowledgments

We thank Zach Heiple, Nick Wegner, Misty Paig-Tran, and Ben Frable for assistance with the recovery and necropsy of the Pacific oarfish specimen. We are grateful to Mark Miller and the FishEvolutionLab for input and criticism during preparation of this manuscript. We thank Pacific Biosciences and the Sanford Consortium Genomics Core Facility (gCore) at UCSD, especially T. Biddle, for assistance with genomic sequencing. Computational analyses were performed thanks to the University of Oklahoma Supercomputing Center for Education & Research (OSCER); we thank Director H. Neeman and System Administrators J. Speckman, T. Ha, and H. Severini for technical support. Further computational support was provided by the San Diego Supercomputer Center (SDSC) and the EXPANSE platform.

## Author contributions

**Conceptualization:** D.A. and F.M.V.

**Methodology:** D.A., F.M.V., J.G.G., W.P., and R.B.R

**Software:** F.M.V., J.G.G., J.Z. and W.P.

**Validation:** D.A., J.G.G., and R.B.R.

**Formal analysis:** D.A., F.M.V., J.G.G., W.P., and R.B.R

**Investigation:** D.A. and F.M.V.

**Resources:** D.A., R.B.R and W.P.

**Data curation:** D.A., F.M.V., J.G.G., J.Z., E.I., and R.B.R

**Writing — original draft:** D.A. and F.M.V.

**Writing — review & editing:** D.A., F.M.V., J.G.G., W.P., E.I., J.Z., and R.B.-R.

**Visualization:** D.A., F.M.V., J.G.G., J.Z., E.I., and R.B.R

**Supervision:** D.A.

**Project administration:** D.A.

**Funding acquisition:** D.A.

## Data availability

Raw PacBio HiFi and Hi-C reads supporting the Pacific oarfish (*Regalecus russellii*) assembly have been deposited at the NCBI Sequence Read Archive under BioProject PRJNA1478024. Processed data, including the Progressive Cactus HAL alignment, CNEE coordinates, PhyloAcc-GT output, MCScanX collinearity files, CAFE5 gene-family tables, and supplementary tables S1–S20, are archived at Zenodo (https://doi.org/10.5281/zenodo.20672095).

## Code availability

All analysis code is archived with a DOI on Zenodo (https://doi.org/10.5281/zenodo.20672095).

## Reporting summary

A completed Nature Portfolio Reporting Summary for ecology, evolution & environmental sciences is provided as part of this submission.

## Supplementary information

Supplementary Information is available for this paper. Figures S1–S22, Tables S1–S21, and Datasets S1–S7 are provided as a separate file and via the Figshare repository linked under Data availability.

## References

1. Legendre, L. J. & Davesne, D. The evolution of mechanisms involved in vertebrate endothermy. Phil. Trans. R. Soc. B 375, 20190136 (2020).

2. Block, B. A. & Finnerty, J. R. Endothermy in fishes: a phylogenetic analysis of constraints, predispositions, and selection pressures. Env. Biol. Fish. 40, 283–302 (1994).

3. Melendez-Vazquez, F. et al. Ecological interactions and genomic innovation fueled the evolution of ray-finned fish endothermy. Sci. Adv. 11, eads8488 (2025).

4. Harding, L. et al. Endothermy makes fishes faster but does not expand their thermal niche. Funct. Ecol. 35, 1951–1959 (2021).

5. Dickson, K. A. & Graham, J. B. Evolution and consequences of endothermy in fishes. Physiol. Biochem. Zool. 77, 998–1018 (2004).

6. Franck, J. P. C., Slight-Simcoe, E. & Wegner, N. C. Endothermy in the smalleye opah (Lampris incognitus): a potential role for the uncoupling protein sarcolipin. Comp. Biochem. Physiol. A 233, 48–52 (2019).

7. Head, G. A., Polymeropoulos, E. T., Oelkrug, R. & Jastroch, M. Editorial: The evolution of endothermy—from patterns to mechanisms. Front. Physiol. 9, 891 (2018).

8. Block, B. A. in The Biochemistry and Molecular Biology of Fishes vol. 1 (eds Hochachka, P. W. & Mommsen, T. P.) 269–311 (Elsevier, 1991).

9. Wegner, N. C., Snodgrass, O. E., Dewar, H. & Hyde, J. R. Whole-body endothermy in a mesopelagic fish, the opah, Lampris guttatus. Science 348, 786–789 (2015).

10. Madigan, D. J. et al. Assessing niche width of endothermic fish from genes to ecosystem. Proc. Natl Acad. Sci. USA 112, 8350–8355 (2015).

11. Block, B. A., Finnerty, J. R., Stewart, A. F. R. & Kidd, J. Evolution of endothermy in fish: mapping physiological traits on a molecular phylogeny. Science 260, 210–214 (1993).

12. Carey, F. G. & Lawson, K. D. Temperature regulation in free-swimming bluefin tuna. Comp. Biochem. Physiol. A 44, 375–392 (1973).

13. Graham, J. B. & Dickson, K. A. The evolution of thunniform locomotion and heat conservation in scombrid fishes: new insights based on the morphology of Allothunnus fallai. Zool. J. Linn. Soc. 129, 419–466 (2000).

14. Camacho García, J. I., et al. Widespread genetic signals of visual system adaptation in deepwater cichlid fishes. Mol. Biol. Evol. 42, msaf001 (2025).

15. Vizueta, J. et al. Adaptive radiation and social evolution of the ants. Cell 188, 1234–1251 (2025).

16. Parey, E. et al. The brittle star genome illuminates the genetic basis of animal appendage regeneration. Nat. Ecol. Evol. 8, 1505–1521 (2024).

17. Morales, A. E. et al. Bat genomes illuminate adaptations to viral tolerance and disease resistance. Nature 638, 449–458 (2025).

18. Marlétaz, F. et al. The hagfish genome and the evolution of vertebrates. Nature 627, 811–820 (2024).

19. Schultz, D. T. et al. Ancient gene linkages support ctenophores as sister to other animals. Nature 618, 110–117 (2023).

20. Shakya, S. B., Edwards, S. V. & Sackton, T. B. Convergent evolution of noncoding elements associated with short tarsus length in birds. BMC Biol. 23, 1–20 (2025).

21. Chen, H. I., Turakhia, Y., Bejerano, G. & Kingsley, D. M. Whole-genome comparisons identify repeated regulatory changes underlying convergent appendage evolution in diverse fish lineages. Mol. Biol. Evol. 40, msad048 (2023).

22. Tang, T. et al. A chromosome-level genome assembly of the reef stonefish (Synanceia verrucosa) provides novel insights into stonustoxin genes. Mol. Biol. Evol. 40, msad046 (2023).

23. Ou, M. et al. Chromosome-level genome assemblies of Channa argus and Channa maculata and comparative analysis of their temperature adaptability. GigaScience 10, giab059 (2021).

24. Xiao, S. et al. Genome of tetraploid fish Schizothorax o’connori provides insights into early re-diploidization and high-altitude adaptation. iScience 23, 101707 (2020).

25. Wu, B. et al. The genomes of two billfishes provide insights into the evolution of endothermy in teleosts. Mol. Biol. Evol. 38, 2413–2427 (2021).

26. Gupta, A., Mirarab, S. & Turakhia, Y. Accurate, scalable, and fully automated inference of species trees from raw genome assemblies using ROADIES. Proc. Natl Acad. Sci. USA 122, e2500553122 (2025).

27. Faircloth, B. C. PHYLUCE is a software package for the analysis of conserved genomic loci. Bioinformatics 32, 786–788 (2016).

28. Carroll, S. B. Evo-devo and an expanding evolutionary synthesis: a genetic theory of morphological evolution. Cell 134, 25–36 (2008).

29. Wittkopp, P. J. & Kalay, G. Cis-regulatory elements: molecular mechanisms and evolutionary processes underlying divergence. Nat. Rev. Genet. 13, 59–69 (2012).

30. Yan, H. et al. PhyloAcc-GT: a Bayesian method for inferring patterns of substitution rate shifts on targeted lineages accounting for gene-tree discordance. Mol. Biol. Evol. 40, msad244 (2023).

31. Armstrong, J. et al. Progressive Cactus is a multiple-genome aligner for the thousand-genome era. Nature 587, 246–251 (2020).

32. Galtier, N. Fine-scale quantification of GC-biased gene conversion intensity in mammals. Peer Community Journal, 1: e17 (2021).

33. Hickey, G., Paten, B., Earl, D., Zerbino, D. & Haussler, D. HAL: a hierarchical format for storing and analyzing multiple genome alignments. Bioinformatics 29, 1341–1342 (2013).

34. Qian, W. & Zhang, J. Genomic evidence for adaptation by gene duplication. Genome Res. 24, 1356–1362 (2014).

35. Wang, Y. et al. MCScanX: a toolkit for detection and evolutionary analysis of gene synteny and collinearity. Nucleic Acids Res. 40, e49 (2012).

36. Mendes, F. K., Vanderpool, D., Fulton, B. & Hahn, M. W. CAFE 5 models variation in evolutionary rates among gene families. Bioinformatics 36, 5516–5518 (2021).

37. Murrell, B. et al. Detecting individual sites subject to episodic diversifying selection. PLOS Genet. 8, e1002764 (2012).

38. Rhie, A. et al. Towards complete and error-free genome assemblies of all vertebrate species. Nature 592, 737–746 (2021).

39. Martin, M. Cutadapt removes adapter sequences from high-throughput sequencing reads. EMBnet J. 17, 10–12 (2011).

40. Rhie, A., Walenz, B. P., Koren, S. & Phillippy, A. M. Merqury: reference-free quality, completeness, and phasing assessment for genome assemblies. Genome Biol. 21, 245 (2020).

41. Ranallo-Benavidez, T. R., Jaron, K. S. & Schatz, M. C. GenomeScope 2.0 and Smudgeplot for reference-free profiling of polyploid genomes. Nat. Commun. 11, 1432 (2020).

42. Cheng, H. et al. Scalable telomere-to-telomere assembly for diploid and polyploid genomes with double graph. Nat. Methods 21, 967–970 (2024).

43. Cheng, H. et al. Haplotype-resolved assembly of diploid genomes without parental data. Nat. Biotechnol. 40, 1332–1335 (2022).

44. Cheng, H., Concepcion, G. T., Feng, X., Zhang, H. & Li, H. Haplotype-resolved de novo assembly using phased assembly graphs with hifiasm. Nat. Methods 18, 170–175 (2021).

45. Guan, D. et al. Identifying and removing haplotypic duplication in primary genome assemblies. Bioinformatics 36, 2896–2898 (2020).

46. Wood, D. E., Lu, J. & Langmead, B. Improved metagenomic analysis with Kraken 2. Genome Biol. 20, 257 (2019).

47. Zhou, C., McCarthy, S. A. & Durbin, R. YaHS: yet another Hi-C scaffolding tool. Bioinformatics 39, btac808 (2023).

48. Pointon, D.-L. B. sanger-tol/curationpretext: CurationPretext pipeline v1.0. Zenodo 10.5281/zenodo.14983419 (2025).

49. Formenti, G. et al. Gfastats: conversion, evaluation and manipulation of genome sequences using assembly graphs. Bioinformatics 38, 4214–4216 (2022).

50. Simão, F. A., Waterhouse, R. M., Ioannidis, P., Kriventseva, E. V. & Zdobnov, E. M. BUSCO: assessing genome assembly and annotation completeness with single-copy orthologs. Bioinformatics 31, 3210–3212 (2015).

51. Storer, J., Hubley, R., Rosen, J., Wheeler, T. J. & Smit, A. F. The Dfam community resource of transposable element families, sequence models, and genome annotations. Mob. DNA 12, 2 (2021).

52. Gabriel, L. et al. BRAKER3: fully automated genome annotation using RNA-seq and protein evidence with GeneMark-ETP, AUGUSTUS and TSEBRA. Genome Res. 34, 769–777 (2024).

53. Katoh, K. & Standley, D. M. MAFFT multiple sequence alignment software version 7: improvements in performance and usability. Mol. Biol. Evol. 30, 772–780 (2013).

54. Minh, B. Q. et al. IQ-TREE 2: new models and efficient methods for phylogenetic inference in the genomic era. Mol. Biol. Evol. 37, 1530–1534 (2020).

55. Wheeler, T. J. & Eddy, S. R. nhmmer: DNA homology search with profile HMMs. Bioinformatics 29, 2487–2489 (2013).

56. Grabherr, M. G., et al. Trinity: reconstructing a full-length transcriptome without a genome from RNA-Seq data. Nat. Biotechnol. 29, 644–652 (2011).

57. Ranwez, V., Douzery, E. J. P., Cambon, C., Chantret, N. & Delsuc, F. MACSE v2: toolkit for the alignment of coding sequences accounting for frameshifts and stop codons. Mol. Biol. Evol. 35, 2582–2584 (2018).

58. Lanfear, R., Frandsen, P. B., Wright, A. M., Senfeld, T. & Calcott, B. PartitionFinder 2: new methods for selecting partitioned models of evolution for molecular and morphological phylogenetic analyses. Mol. Biol. Evol. 34, 772–773 (2017).

59. Kalyaanamoorthy, S., Minh, B. Q., Wong, T. K. F., von Haeseler, A. & Jermiin, L. S. ModelFinder: fast model selection for accurate phylogenetic estimates. Nat. Methods 14, 587–589 (2017).

60. Pennell, M. W. et al. geiger v2.0: an expanded suite of methods for fitting macroevolutionary models to phylogenetic trees. Bioinformatics 30, 2216–2218 (2014).

61. Bandi, V. & Gutwin, C. SynVisio: an interactive multi-scale visualization tool for genome synteny. https://synvisio.github.io (2020).

62. Eastman, J. M., Harmon, L. J. & Tank, D. C. Congruification: support for time scaling large phylogenetic trees. Methods Ecol. Evol. 4, 688–691 (2013).

63. Emms, D. M. & Kelly, S. OrthoFinder: phylogenetic orthology inference for comparative genomics. Genome Biol. 20, 238 (2019).

64. Wu, T. et al. clusterProfiler 4.0: a universal enrichment tool for interpreting omics data. Innovation 2, 100141 (2021).

65. Cantalapiedra, C. P., Hernández-Plaza, A., Letunic, I., Bork, P. & Huerta-Cepas, J. eggNOG-mapper v2: functional annotation, orthology assignments, and domain prediction at the metagenomic scale. Mol. Biol. Evol. 38, 5825–5829 (2021).

66. Gene Ontology Consortium. The Gene Ontology knowledgebase in 2023. Genetics 224, iyad031 (2023).

67. Supek, F., Bošnjak, M., Škunca, N. & Šmuc, T. REVIGO summarizes and visualizes long lists of gene ontology terms. PLOS ONE 6, e21800 (2011).

68. Kosakovsky Pond, S. L., et al. HyPhy 2.5—a customizable platform for evolutionary hypothesis testing using phylogenies. Mol. Biol. Evol. 37, 295–299 (2020).

69. Hubisz, M. J., Pollard, K. S. & Siepel, A. PHAST and Rphast: phylogenetic analysis with space/time models. Brief. Bioinform. 12, 41–51 (2011).

70. Zhang, C., Rabiee, M., Sayyari, E. & Mirarab, S. ASTRAL-III: polynomial time species tree reconstruction from partially resolved gene trees. BMC Bioinformatics 19, 153 (2018).

71. Sackton, T. B. et al. Convergent regulatory evolution and loss of flight in paleognathous birds. Science 364, 74–78 (2019).

72. Baranasic, D. et al. Multiomic atlas with functional stratification and developmental dynamics of zebrafish cis-regulatory elements. Nat. Genet. 54, 1037–1050 (2022).

73. Ciezarek, A., Gardner, L., Savolainen, V. & Block, B. A. Skeletal muscle and cardiac transcriptomics of a regionally endothermic fish, the Pacific bluefin tuna, Thunnus orientalis. BMC Genomics 21, 642 (2020).

74. Wang, X. et al. Genomic basis of evolutionary adaptation in a warm-blooded fish. The Innovation 3, 100185 (2022).

75. Avsec, Ž., et al. Advancing regulatory variant effect prediction with AlphaGenome. Nature 10.1038/s41586-025-10014-0 (2026).

76. Wainwright, P. C., Alfaro, M. E., Bolnick, D. I. & Hulsey, C. D. Many-to-one mapping of form to function: a general principle in organismal design? Integr. Comp. Biol. 45, 256–262 (2005).

77. Stoltzfus, A. & Norris, R. W. On the causes of evolutionary transition:transversion bias. Mol. Biol. Evol. 33, 595–602 (2016).

78. Reams, A. B. & Neidle, E. L. Selection for gene clustering by tandem duplication. Annu. Rev. Microbiol. 58, 119–142 (2004).

