## Supplementary Materials for "Independent origins of fish endothermy converge on a developmental regulatory signature"

**Supplementary Figures S1–S22 (this file)**

**Supplementary Tables S1–S20 (separate Excel workbook; see Data Availability)**

**Supplementary Tables S21 (this file)**

**Supplementary Methods (this file)**

**Supplementary Datasets S1–S7** (separate archive; see Data Availability)

### Supplementary Methods

#### Targeted MEME re-analysis of prior candidate genes.

Site-specific (codon) signals of episodic diversifying selection across EET2 and EET3 were analyzed with the Mixed Effects Model of Evolution (MEME) in HyPhy v2.5. Analyses were restricted to dcaf6 and carnmt1, nominated by Melendez-Vazquez et al. (ref. 3) as candidates with convergent positive selection across endothermic vertebrate scenarios (carnmt1 in regional and full-body scenarios; dcaf6 in all-endotherm and eye/brain scenarios). The pruned ROADIES phylogram was used as the guide tree. Endothermic species were assigned as foreground and ectothermic species as background. L. incognitus was excluded because the full-body convergent signal in ref. 3 was dominated by non-fish endotherms (marine mammals and penguins), so the targeted MEME re-analysis was focused on the EET2 and EET3 ray-finned fish lineages where ref. 3 reported scenario-specific signals. Per-codon results are reported in Tables S12 and S13 and visualized in Figs. S14 and S15.

### Supplementary Figures

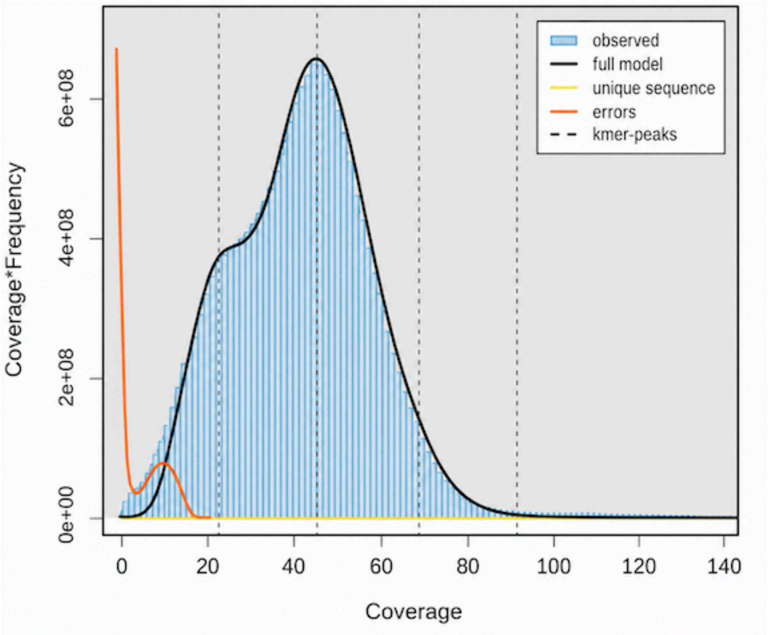


**Fig. S1. GenomeScope 2.0 profile of *Regalecus russellii*.** K-mer spectrum of the *Regalecus russellii* Illumina reads generated using GenomeScope 2.0. The estimated genome size is ~579 Mb with 85.9% unique sequence, 1.02% heterozygosity, and 1.45% estimated duplication. K-mer coverage was modelled at 22.6× with an error rate of 0.166% using a k-mer length of 31. The final assembly (746 Mb) exceeds the k-mer-based estimate (~579 Mb) by ~22%, consistent with MDA-induced k-mer skew and retained haplotypic duplication (see Methods); coverage uniformity across chromosome-scale scaffolds (median ratio 0.95–1.05) and the 5% BUSCO duplication rate support this interpretation.

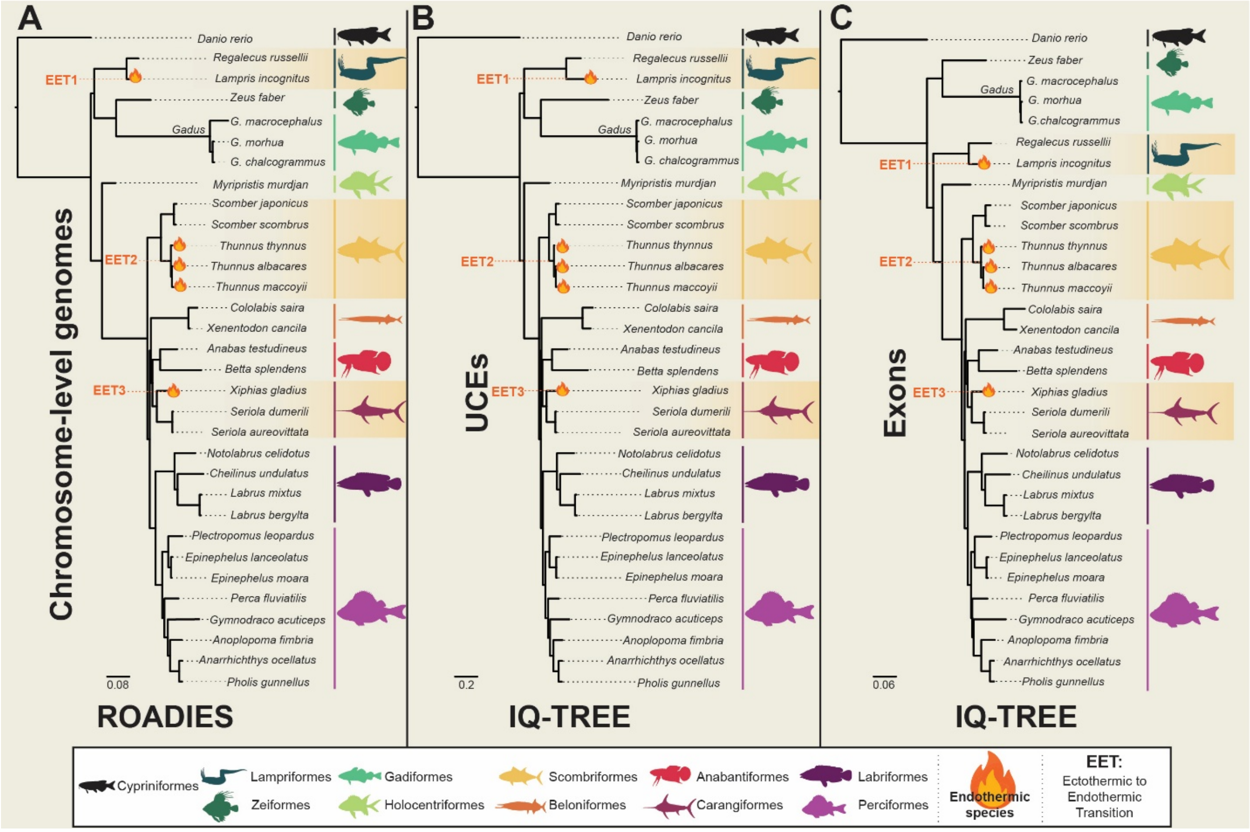


**Fig. S2. Phylogenetic relationships inferred from three independent genomic datasets.** Side-by-side species trees inferred from (A) chromosome-level genome assemblies using ROADIES (~32,000 single- and multi-copy gene trees), (B) 1,095 ultraconserved elements (UCEs) under IQ-TREE with the MIX model, and (C) 1,105 single-copy nuclear exons under IQ-TREE with PartitionFinder and ModelFinder. Trees are time-calibrated where indicated; branch support is shown as bootstrap or local posterior probability. Topological congruence across datasets is restricted to opahs, tunas, and billfishes; discordance is confined to deeper acanthomorph nodes and does not affect the identity of the ectothermic–endothermic sister lineages used in downstream contrasts. Newick files are provided in Dataset S1.

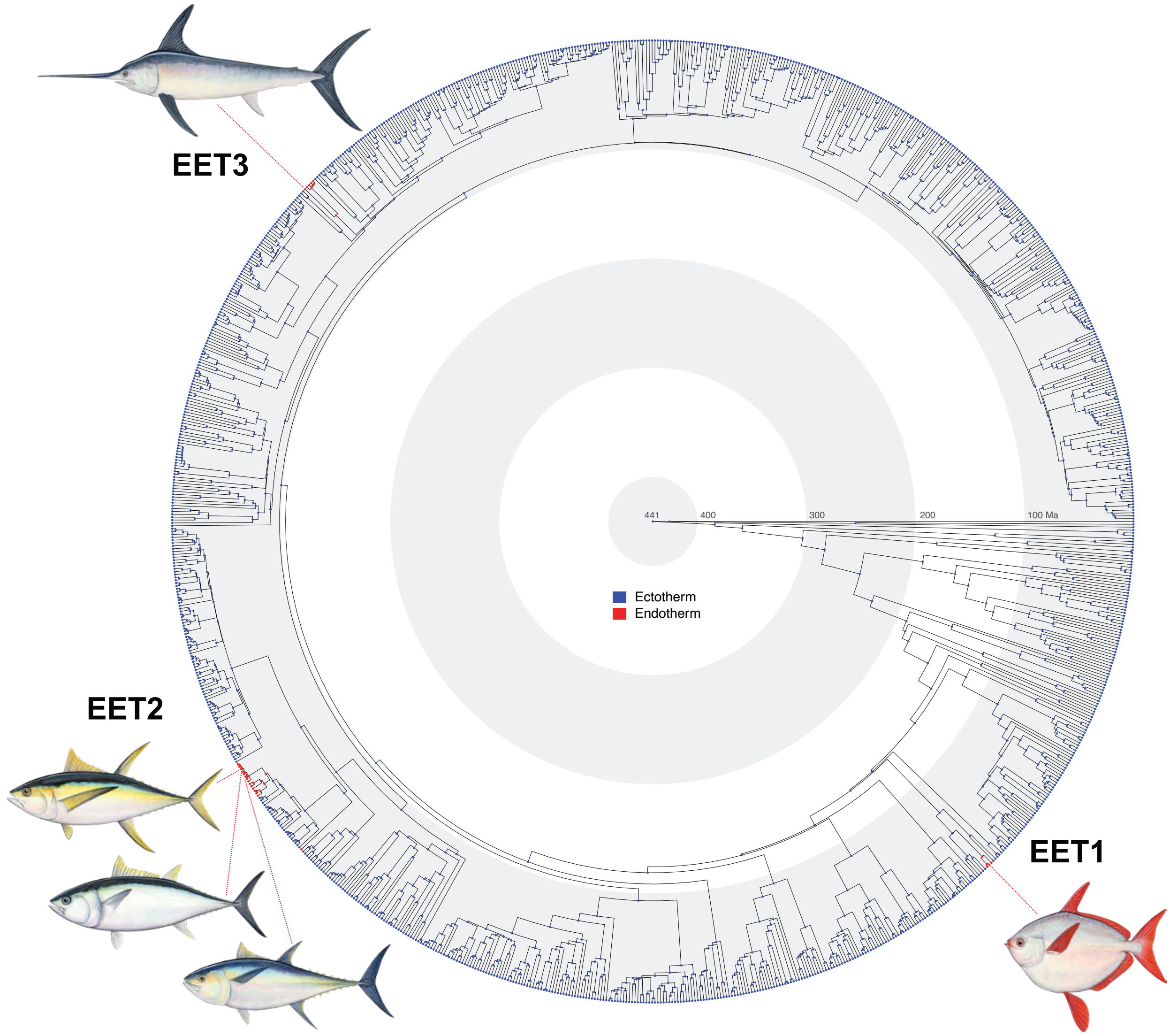


**Fig. S3. Ancestral state reconstruction of endothermy across 1,051 ray-finned fishes.** Phylogenomic tree replotted from Melendez-Vazquez et al. 2025 (ref. 3). Stochastic character mapping of the presence (red) or absence (blue) of endothermy under an equal-rates model demonstrates strong support for endothermy at the nodes subtending the three focal endothermic lineages and ectothermy at all other nodes. Blue-to-red gradient arrows indicate branches along which the ectotherm-to-endotherm transition occurred. This reconstruction is used to infer transitions rather than the 32-taxon tree because of its superior sampling, which more accurately reflects endothermy as a rare trait. See also HiSSE-based reconstructions in Dataset S7.

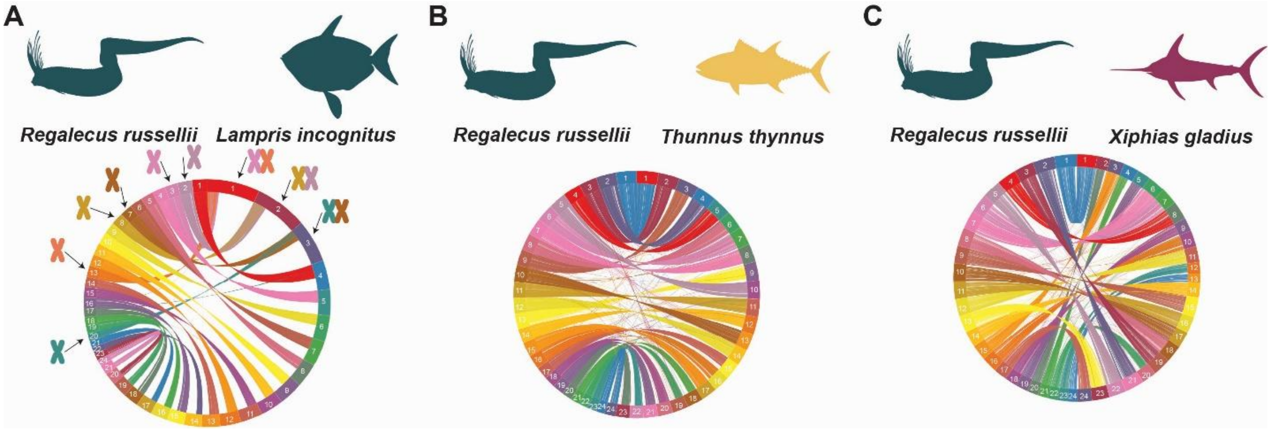


**Fig. S4. Pairwise chromosomal synteny between *Regalecus russellii* and representative endothermic species from EET2 and EET3.** Synteny maps comparing chromosome-level assemblies of *Regalecus russellii* to (A) *Lampris incognitus* (whole-body endotherm, EET1), (B) *Thunnus thynnus* (regional endotherm, EET2), and (C) *Xiphias gladius* (cranial endotherm, EET3). Colors denote homologous chromosomes; connecting ribbons indicate conserved collinear blocks.

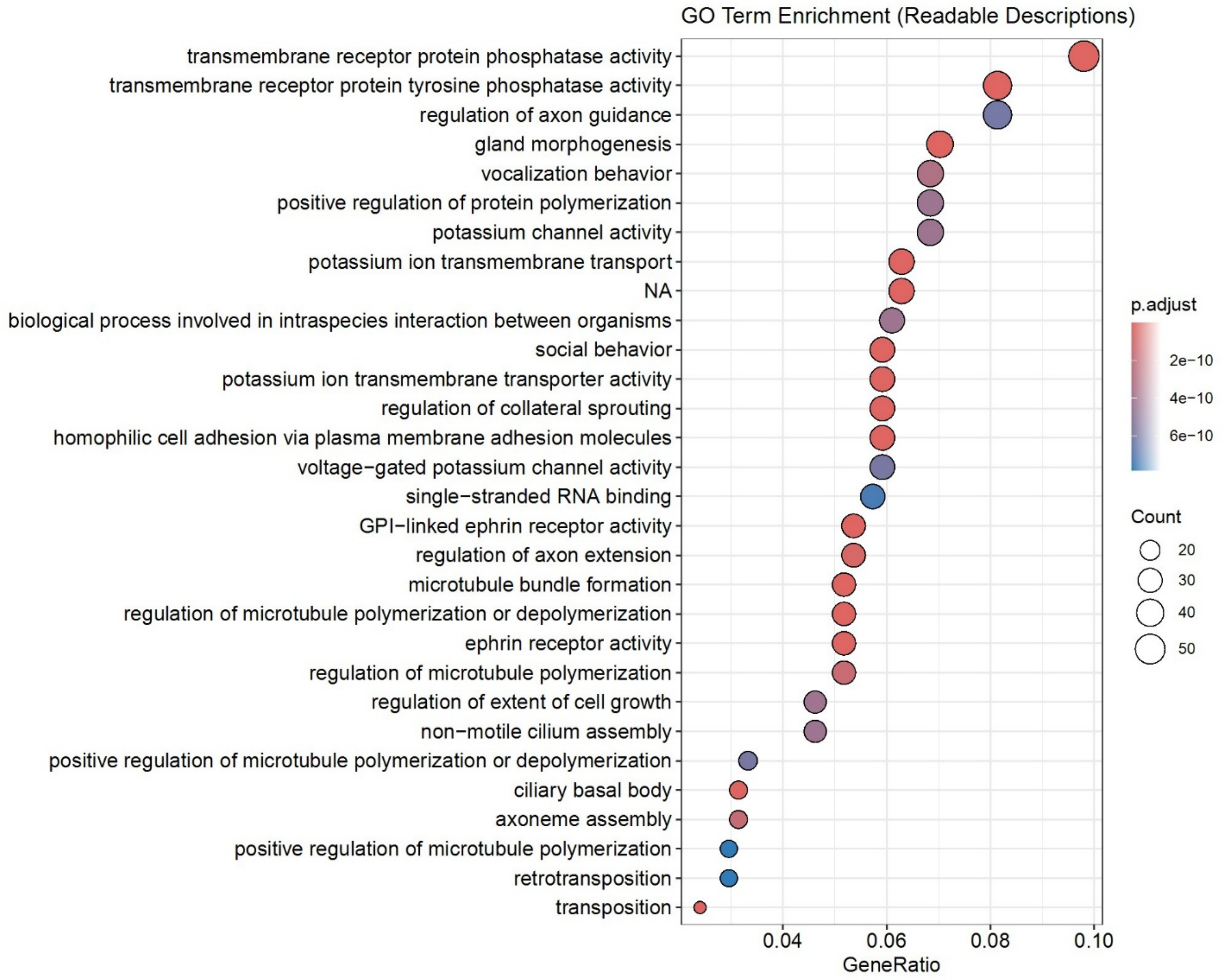


**Fig. S5. Top 30 enriched GO Biological Process categories in *Regalecus russellii***. Bubble plot showing the most significantly enriched GO terms among genes under expansion in *Regalecus russellii*. Gene ratios represent the proportion of expanded genes assigned to each GO category. Bubble color indicates adjusted p-values; bubble size reflects the number of genes associated with each term.

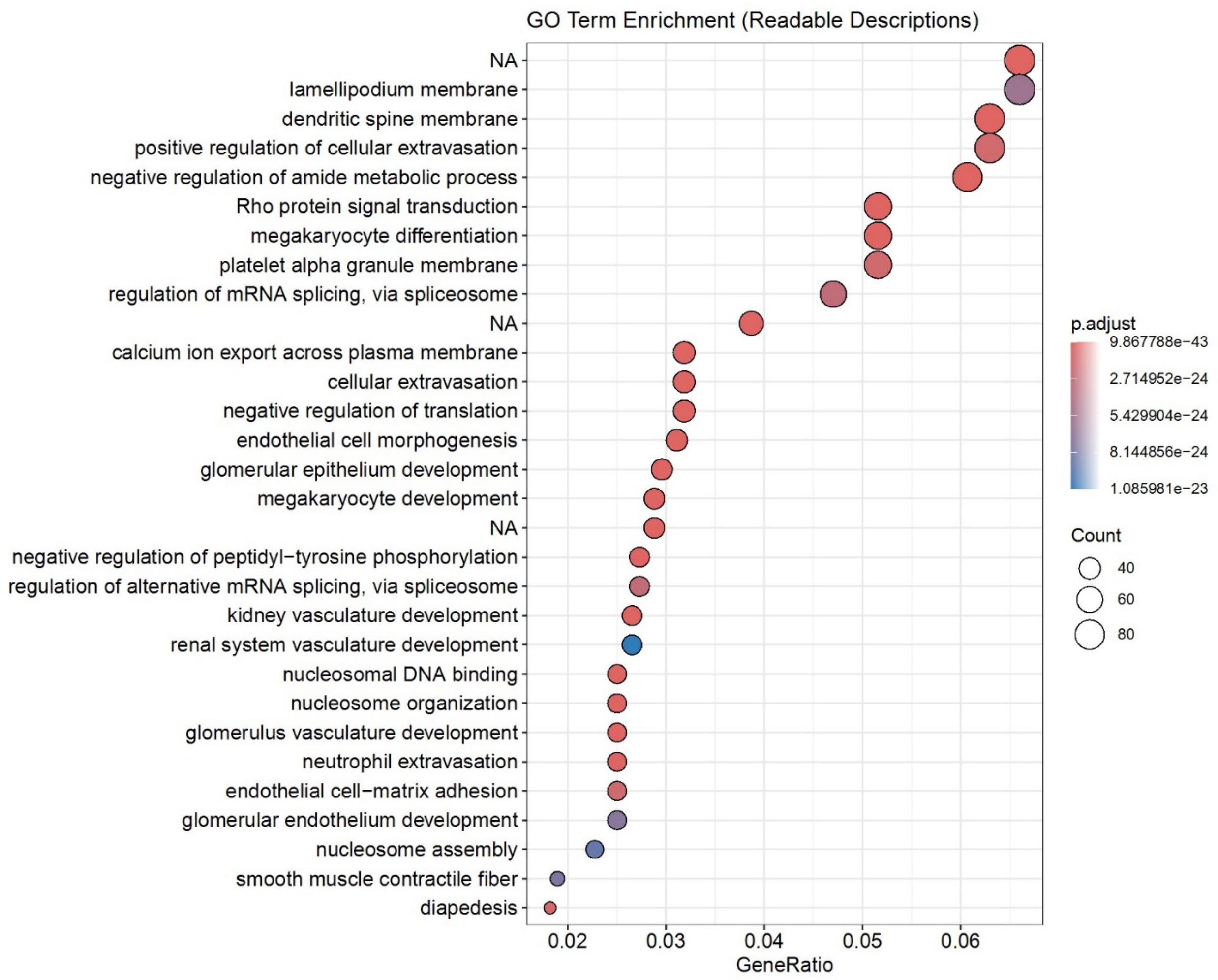


**Fig. S6. Top 30 enriched GO Biological Process categories in *Lampris incognitus***. Bubble plot showing the most significantly enriched GO terms among genes under expansion in *Lampris incognitus*. Gene ratios represent the proportion of expanded genes assigned to each GO category. Bubble color indicates adjusted p-values; bubble size reflects the number of genes associated with each term.

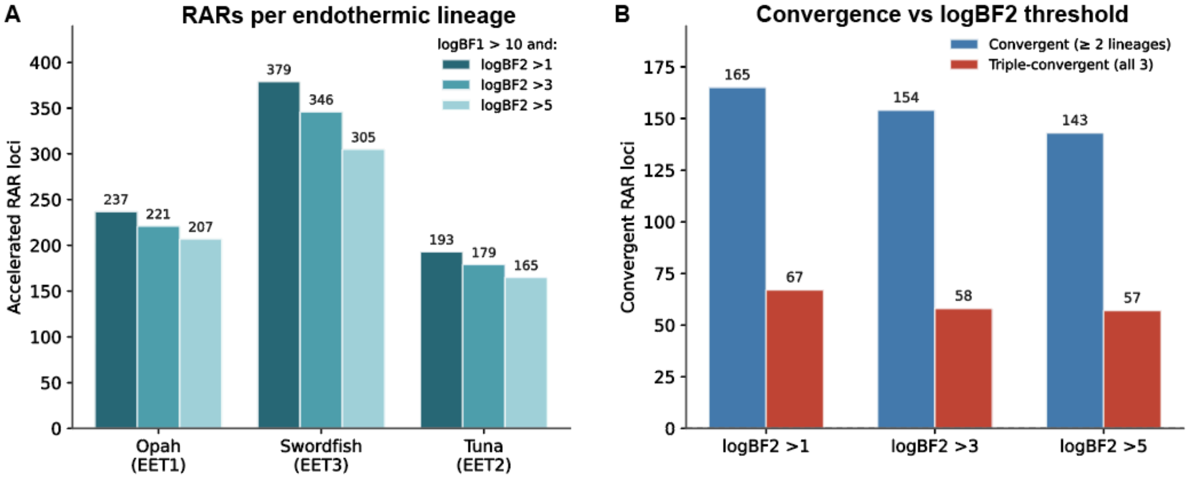


**Fig. S7. Sensitivity of PhyloAcc-GT convergence results to logBF2 threshold.** (A) Number of EARs per endothermic lineage and (B) numbers of pairwise and triple-convergent EARs at logBF1 > 10 paired with three logBF2 cutoffs (>1, the threshold used in the main analysis; >3, substantial support per Kass & Raftery; >5, strong support). At progressively stricter cutoffs the total EAR set reduces from 577 (logBF2 > 1) to 534 (> 3) and 477 (> 5), while the triple-convergent count (67, 58, and 57, respectively) remains substantially in excess of the permutation null (median = 0; empirical $P < 10^{-4}$ at all cutoffs). The convergence signal is robust to the choice of logBF2 threshold.

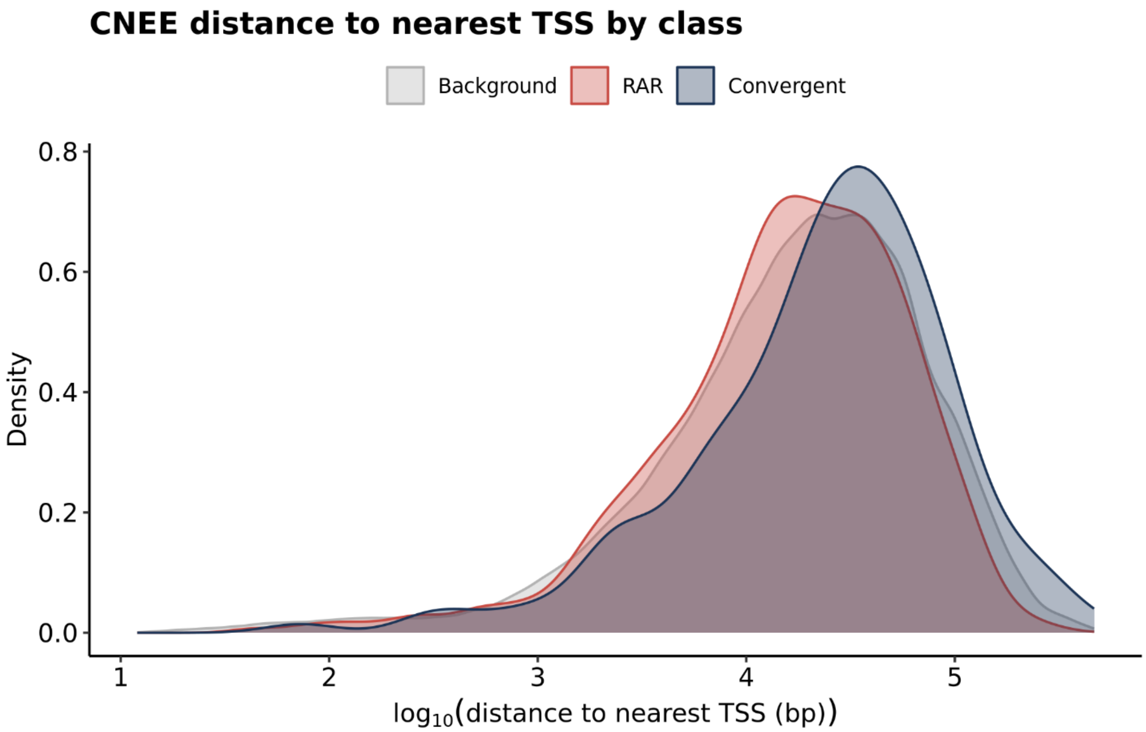


**Fig. S8. Distance of accelerated conserved noncoding elements to nearest transcription start site.** Density distributions of $\log_{10}$-transformed distances between conserved noncoding elements (CNEEs) and the nearest transcription start site (TSS) for background CNEEs (gray), all endothermic accelerated regulatory regions (EARs; red), and convergent EARs shared across endothermic lineages (blue). Convergent EARs sit at slightly greater median distances from TSS than lineage-specific EARs ($\log_{10}$ distance ≈ 4.5 vs. 4.2), consistent with action through distal enhancers.

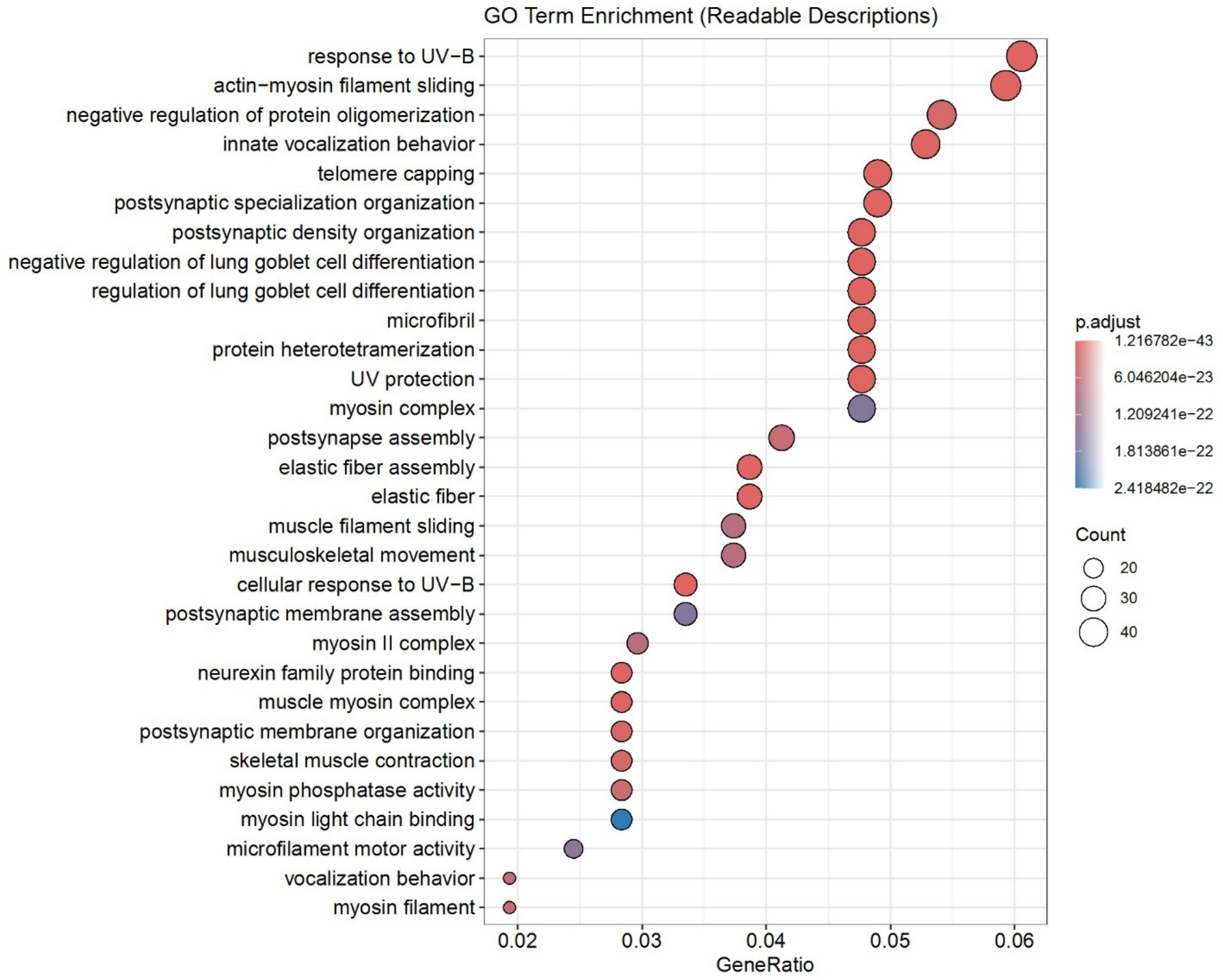


**Fig. S9. Top 30 enriched Gene Ontology (GO) categories in *Thunnus thynnus***. Bubble plot showing the top 30 significantly enriched GO terms among expanded gene families in the regional endotherm *T. thynnus*. Gene ratios represent the proportion of expanded genes assigned to each GO category. Bubble color indicates adjusted p-values; bubble size reflects the number of genes associated with each term.

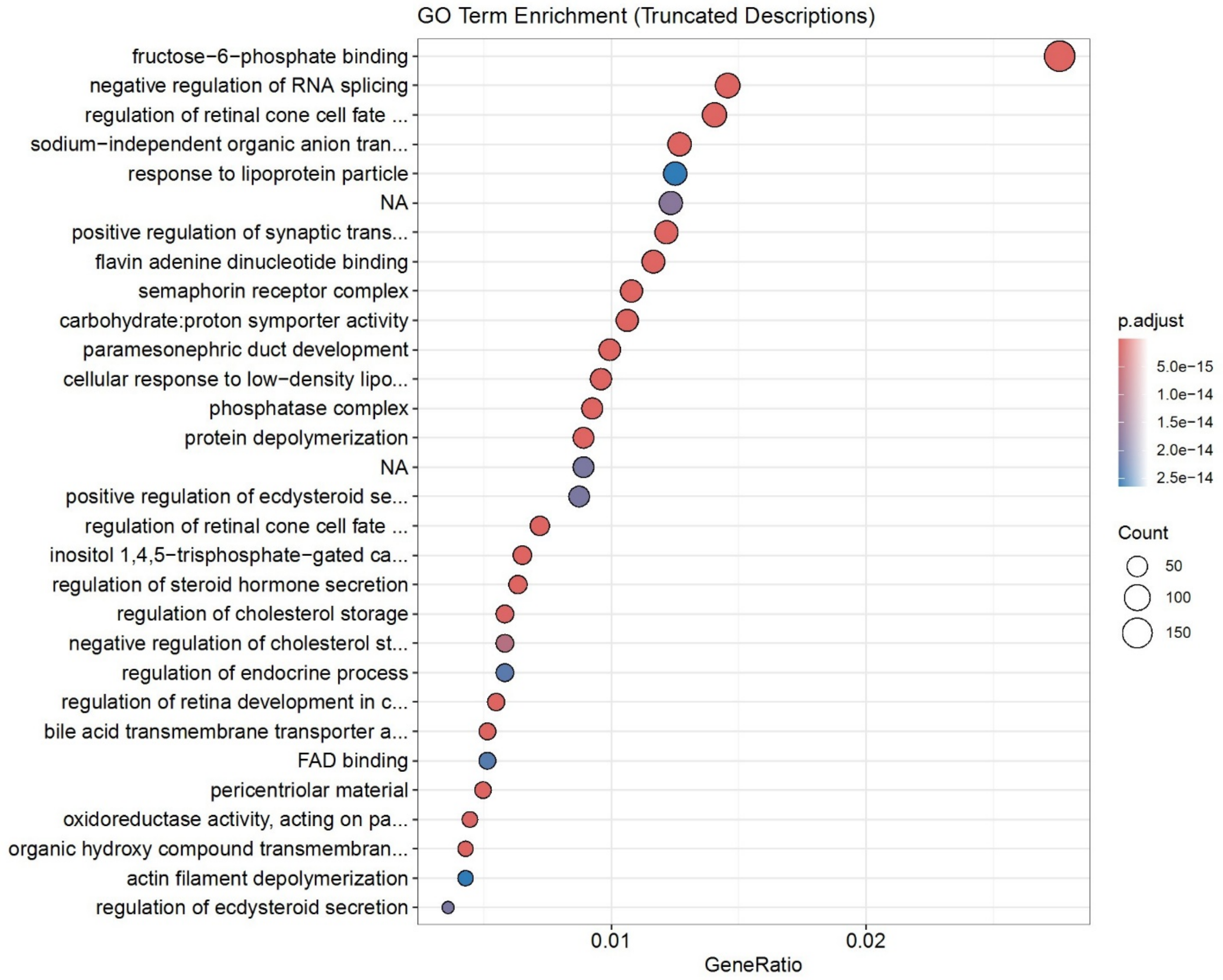


**Fig. S10. Top 30 enriched Gene Ontology (GO) categories in *Xiphias gladius***. Bubble plot showing the top 30 significantly enriched GO terms among expanded gene families in the cranial endotherm *X. gladius*. Gene ratios represent the proportion of expanded genes assigned to each GO category. Bubble color indicates adjusted p-values; bubble size reflects the number of genes associated with each term.

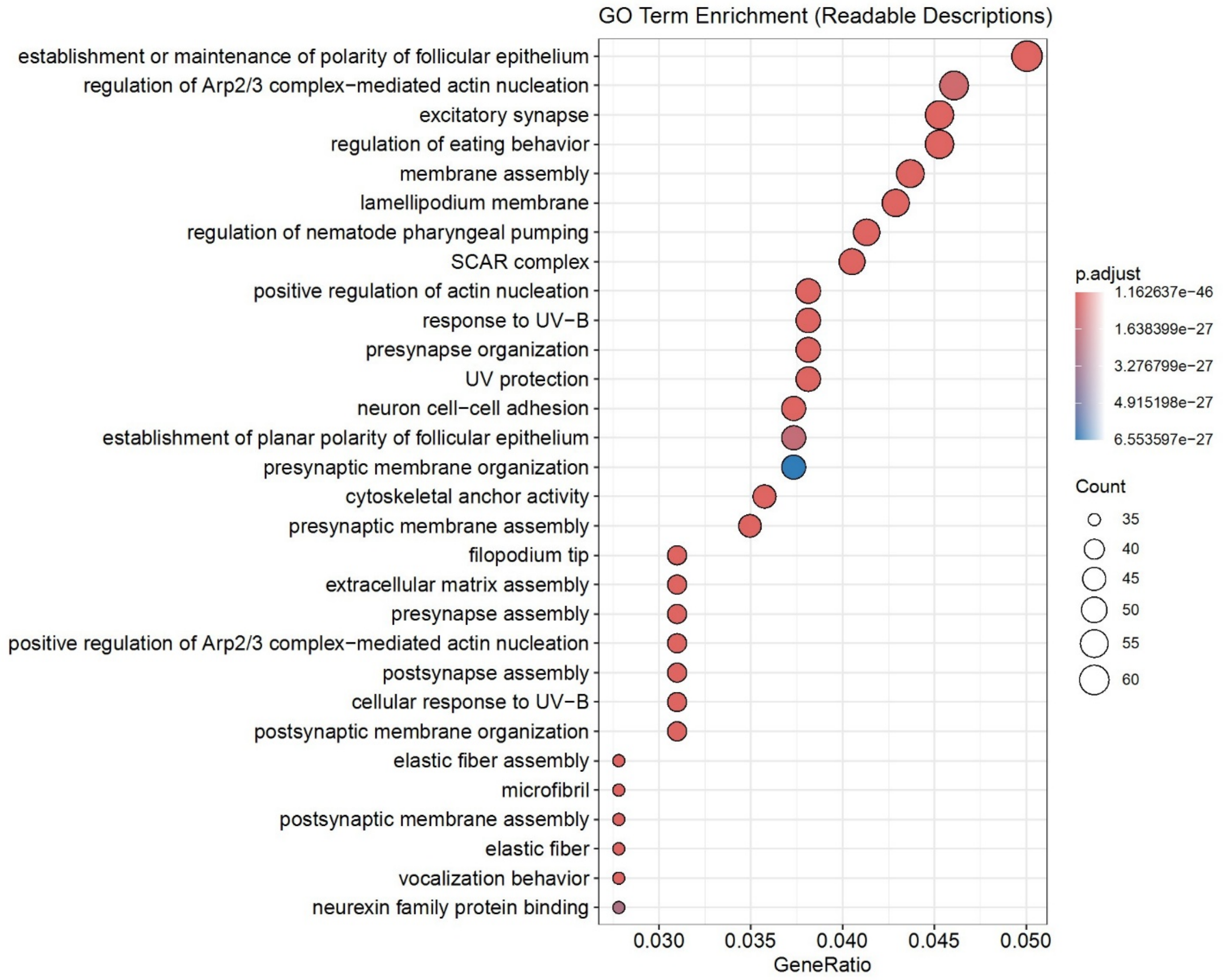


**Fig. S11. Top 30 enriched Gene Ontology (GO) categories in *Scomber japonicus***. Bubble plot showing the top 30 significantly enriched GO terms among expanded gene families in the ectothermic sister of EET2, *S. japonicus*. Gene ratios represent the proportion of expanded genes assigned to each GO category. Bubble color indicates adjusted p-values; bubble size reflects the number of genes associated with each term.

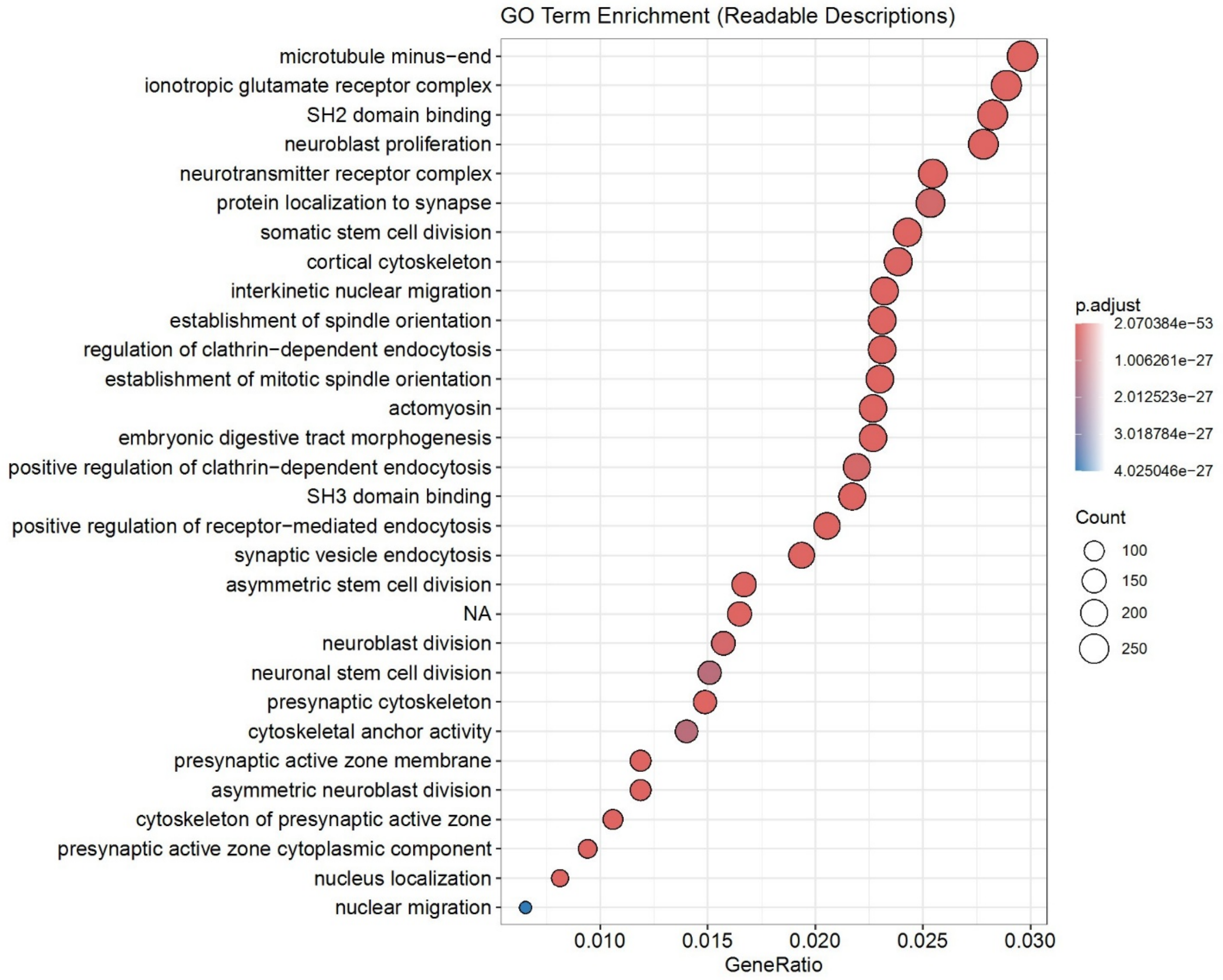


**Fig. S12. Top 30 enriched Gene Ontology (GO) categories in *Seriola aureovittata***. Bubble plot showing the top 30 significantly enriched GO terms among expanded gene families in the ectothermic sister of EET3, *S. aureovittata*. Gene ratios represent the proportion of expanded genes assigned to each GO category. Bubble color indicates adjusted p-values; bubble size reflects the number of genes associated with each term.

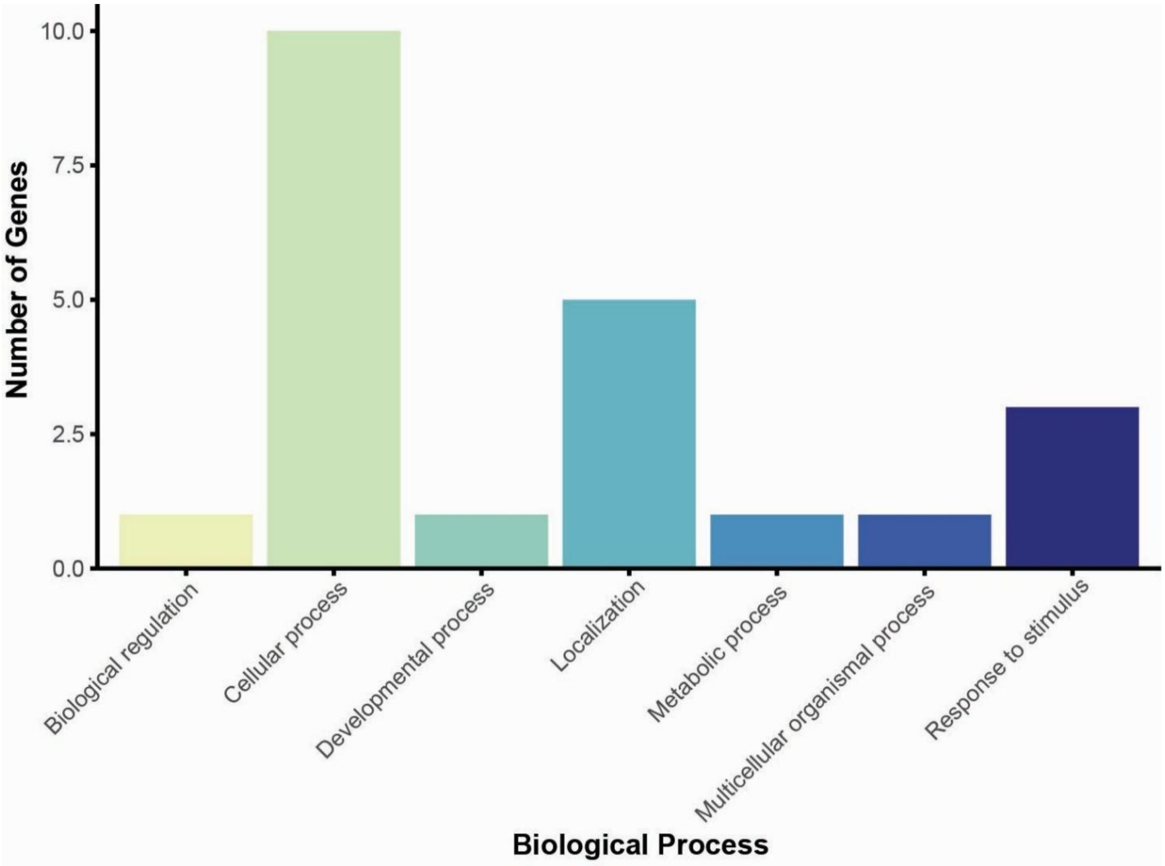


**Fig. S13. PANTHER GO-slim Biological Process classification of gene families exclusively expanded in endothermic fishes.** Bar chart showing the distribution of biological process categories for the 24 gene families ~~convergently~~ expanded across the three independent endothermic transitions (EET1–EET3). Categories include cytoskeletal remodeling, photoreceptor and optic-nerve development, and immune or stress-related processes.

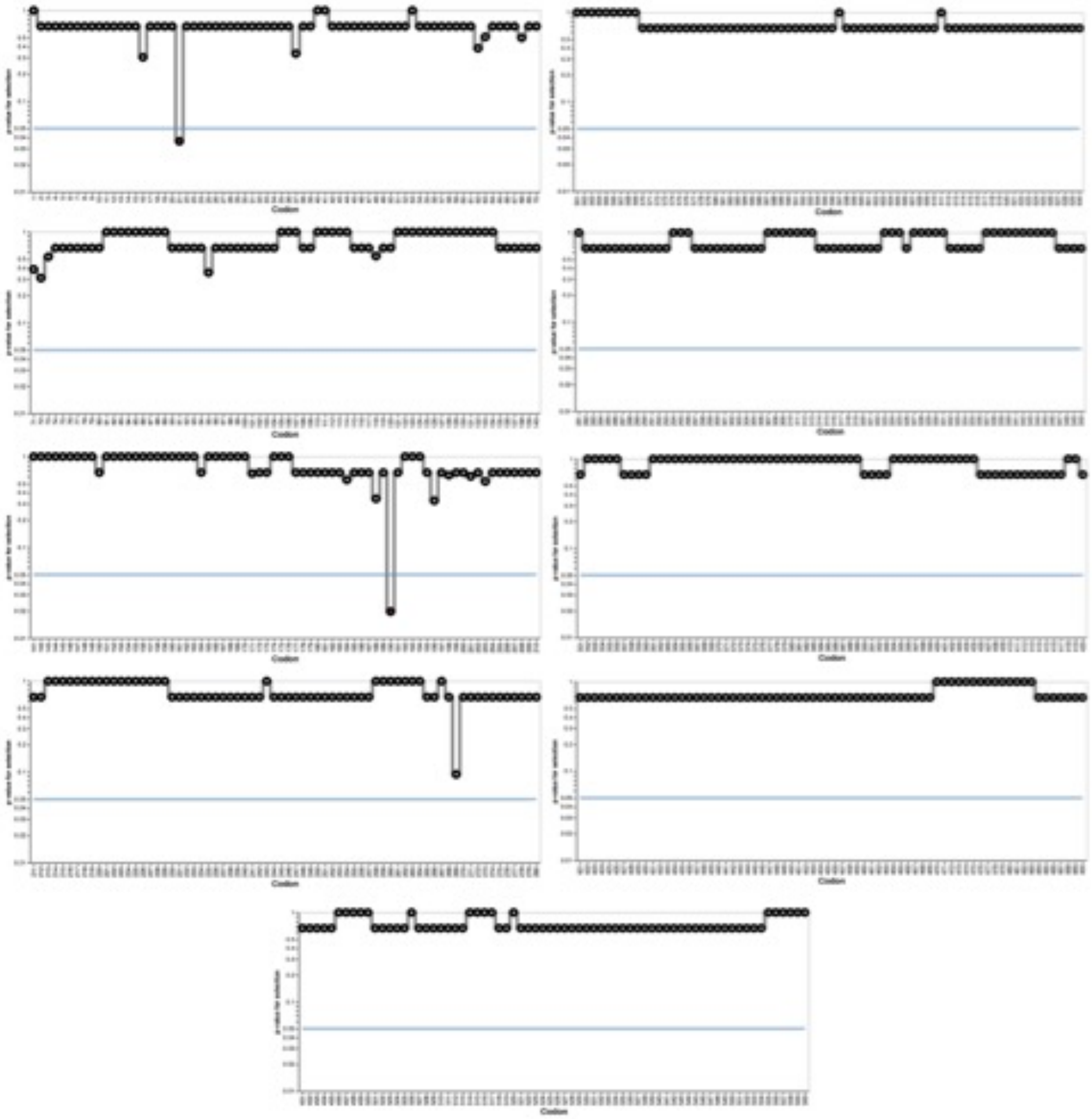

**Fig. S14. Signatures of episodic selection for *carnmt1*.** Codon-level evidence of episodic positive selection in endothermic lineages identified using the Mixed Effects Model of Evolution (MEME), with endothermic representatives used as foreground in EET2 and EET3. Codon positions for *carnmt1* span sites 1–560. The horizontal dashed line indicates the significance threshold ($p < 0.05$); significant sites are marked in red.

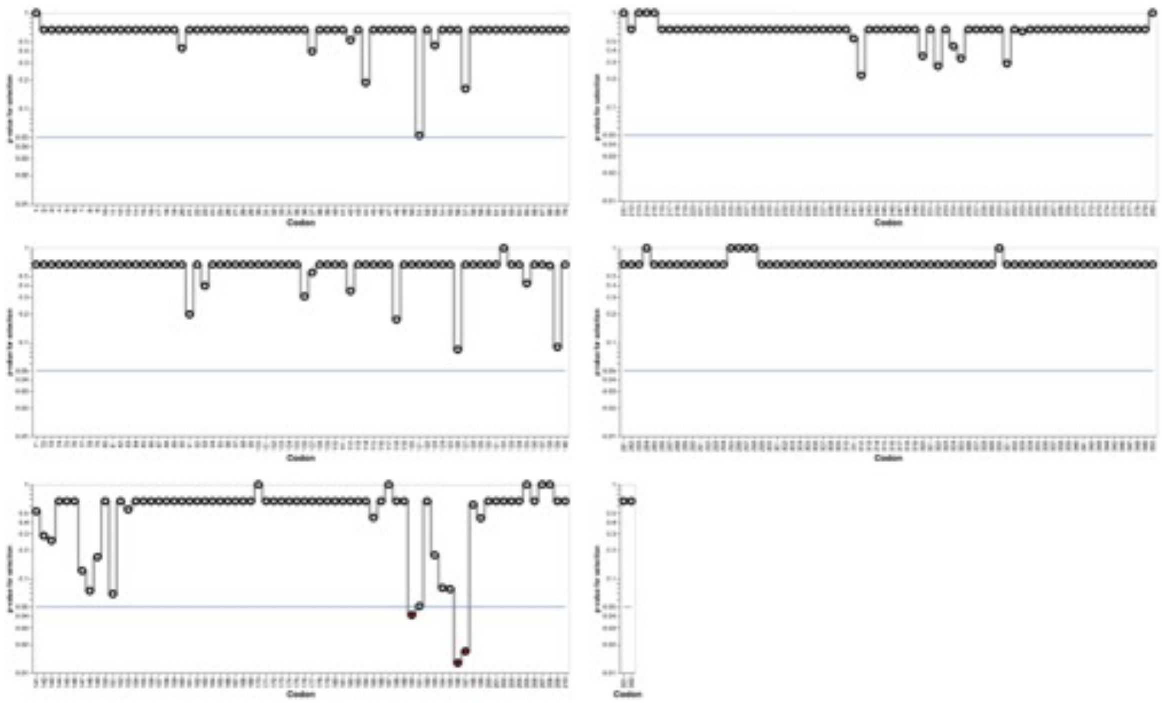

**Fig. S15. Signatures of episodic selection for *dcaf6***. Codon-level evidence of episodic positive selection in endothermic lineages identified using the Mixed Effects Model of Evolution (MEME), with endothermic representatives used as foreground in EET2 and EET3. Codon positions for *dcaf6* span sites 1–352. The horizontal dashed line indicates the significance threshold ($p < 0.05$); significant sites are marked in red.

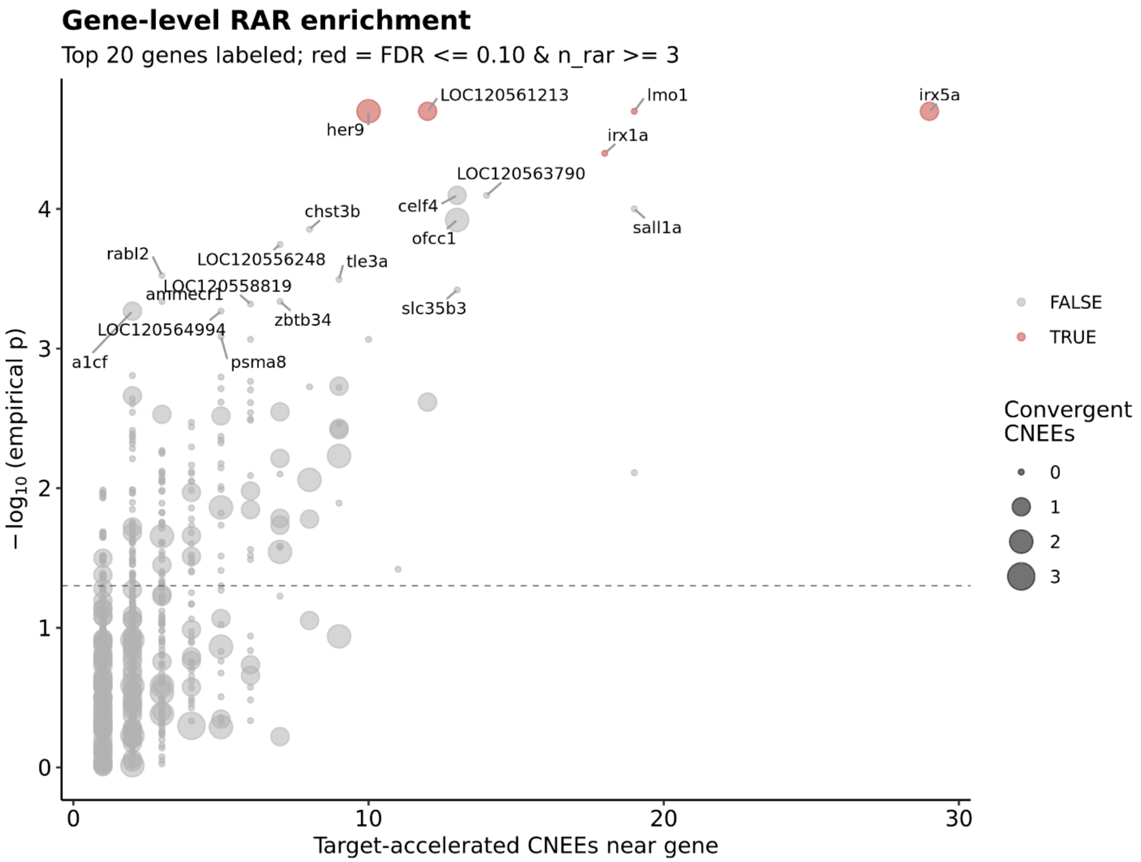


**Fig. S16. Gene-level enrichment of endothermic accelerated conserved noncoding elements (EARs) across endothermic lineages, extended view.** Each point represents one annotated gene; the x-axis shows the number of target-accelerated CNEEs near the gene and the y-axis shows the empirical significance ($-\log_{10}$ p). Point size denotes the number of convergent CNEEs associated with each gene. Genes highlighted in red passed the enrichment threshold (FDR $\leq$ 0.05 with EAR $\geq$ 3). The dashed horizontal line marks the nominal significance threshold (p = 0.05). This figure complements main-text Fig. 2E by showing the full background distribution of CNEE-proximal genes.

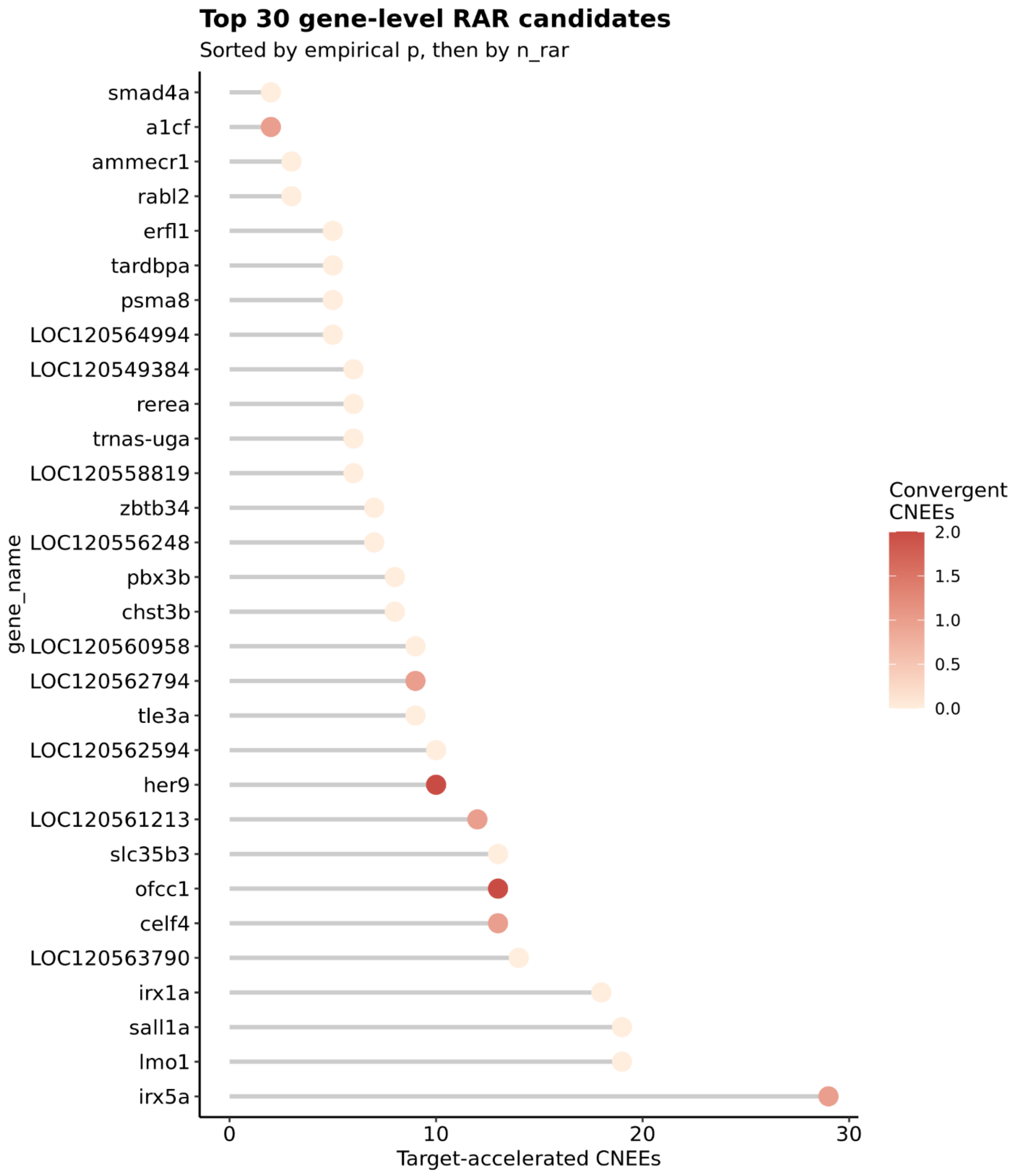


**Fig. S17. Top 30 gene-level candidates associated with endothermic accelerated conserved noncoding elements (EARs).** Genes ranked by empirical significance and number of target-accelerated CNEEs. Each point represents one annotated gene, with the x-axis showing the number of accelerated CNEEs associated with that locus. Point color denotes the number of convergent CNEEs shared across endothermic lineages.

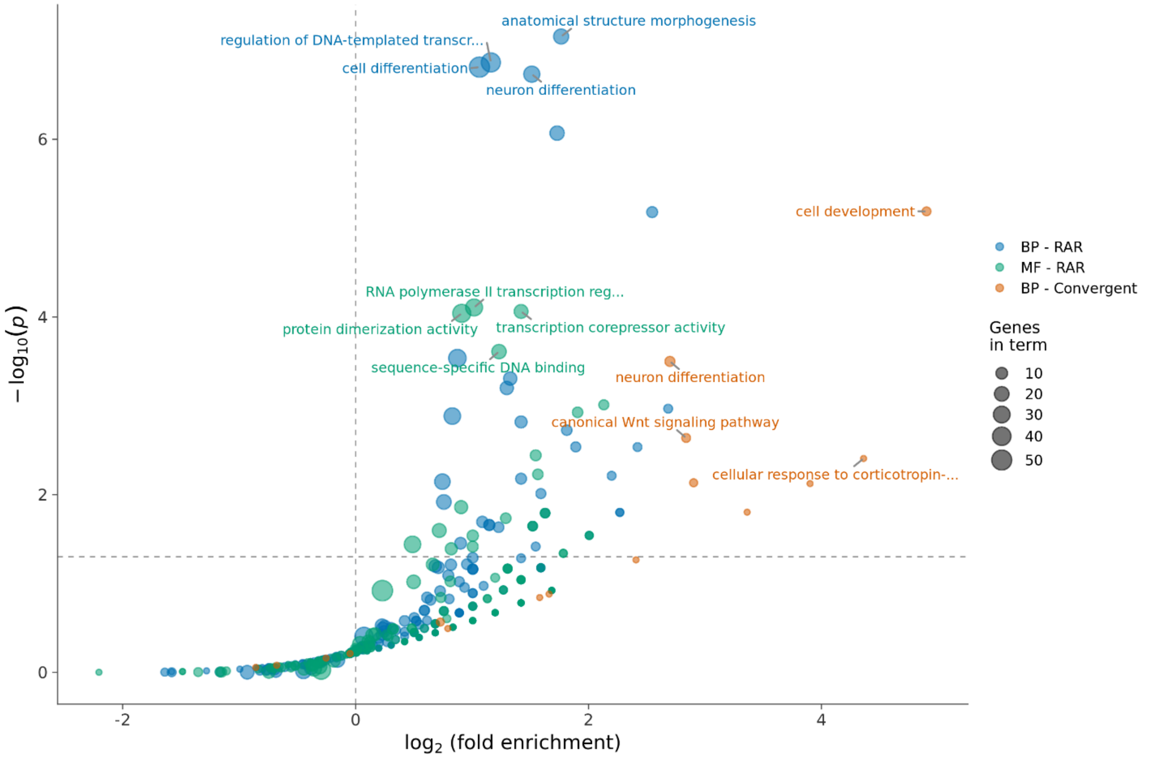


**Fig. S18. Gene Ontology (GO) enrichment associated with endothermic accelerated conserved noncoding elements (EARs), extended view.** Each point represents one enriched GO term, plotted by fold enrichment (x-axis) and statistical significance ($-\log_{10}$ p; y-axis). Point size denotes the number of genes associated with each term. Colors distinguish biological-process terms associated with all EARs (BP–EAR), molecular-function terms associated with all EARs (MF–EAR), and biological-process terms associated with the convergent subset of EARs shared across endothermic lineages (BP–Convergent). Dashed lines indicate nominal enrichment and significance thresholds.

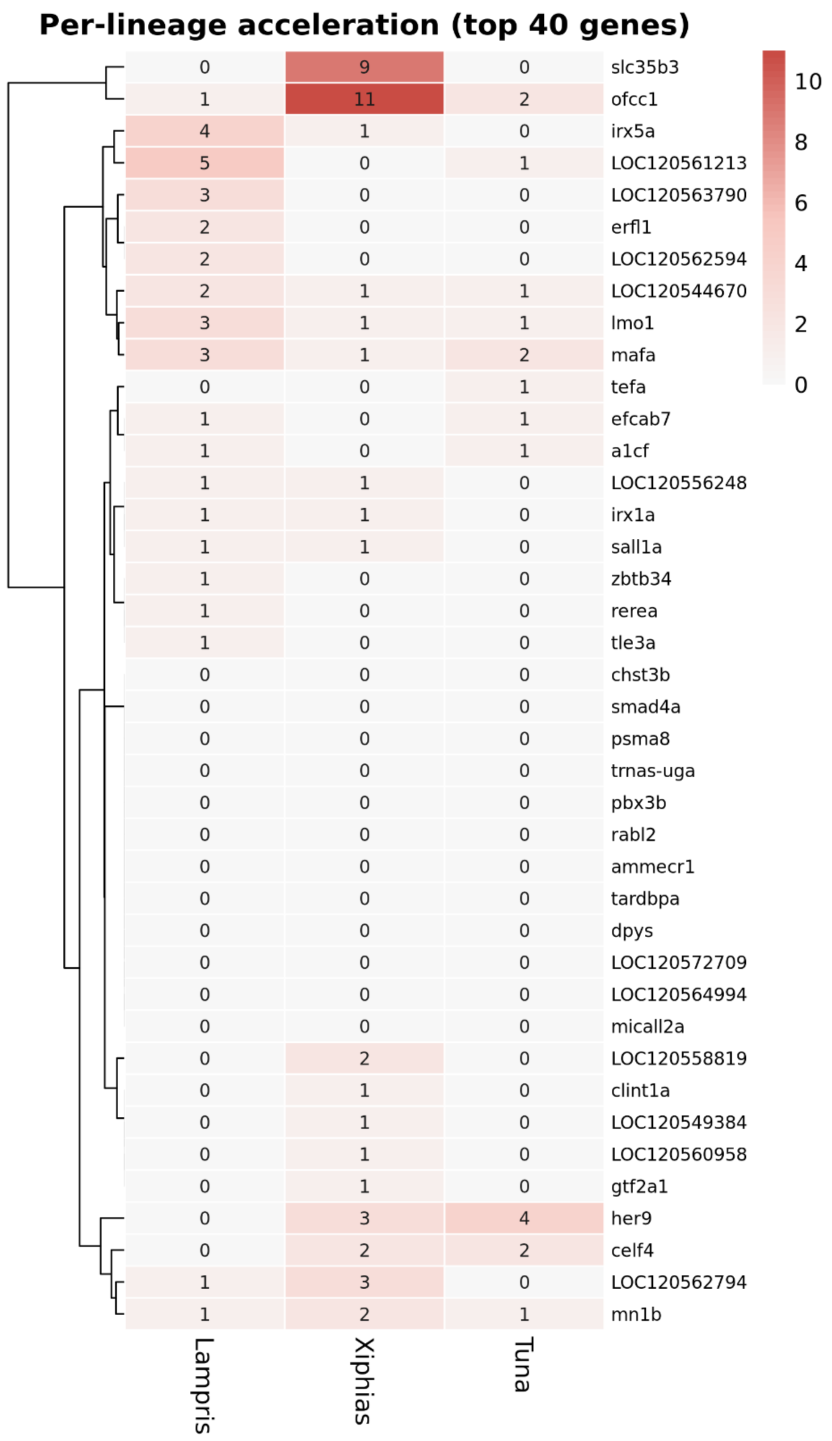


**Fig. S19. Per-lineage distribution of accelerated regulatory elements across top candidate genes.** Heatmap showing the distribution of target-accelerated CNEEs (EARs) associated with the top 40 candidate genes across the three independent endothermic lineages: *Lampris* (opah), *Xiphias* (swordfish), and tunas. Cell values indicate the number of accelerated CNEEs associated with each gene in each lineage; darker shading represents higher counts.

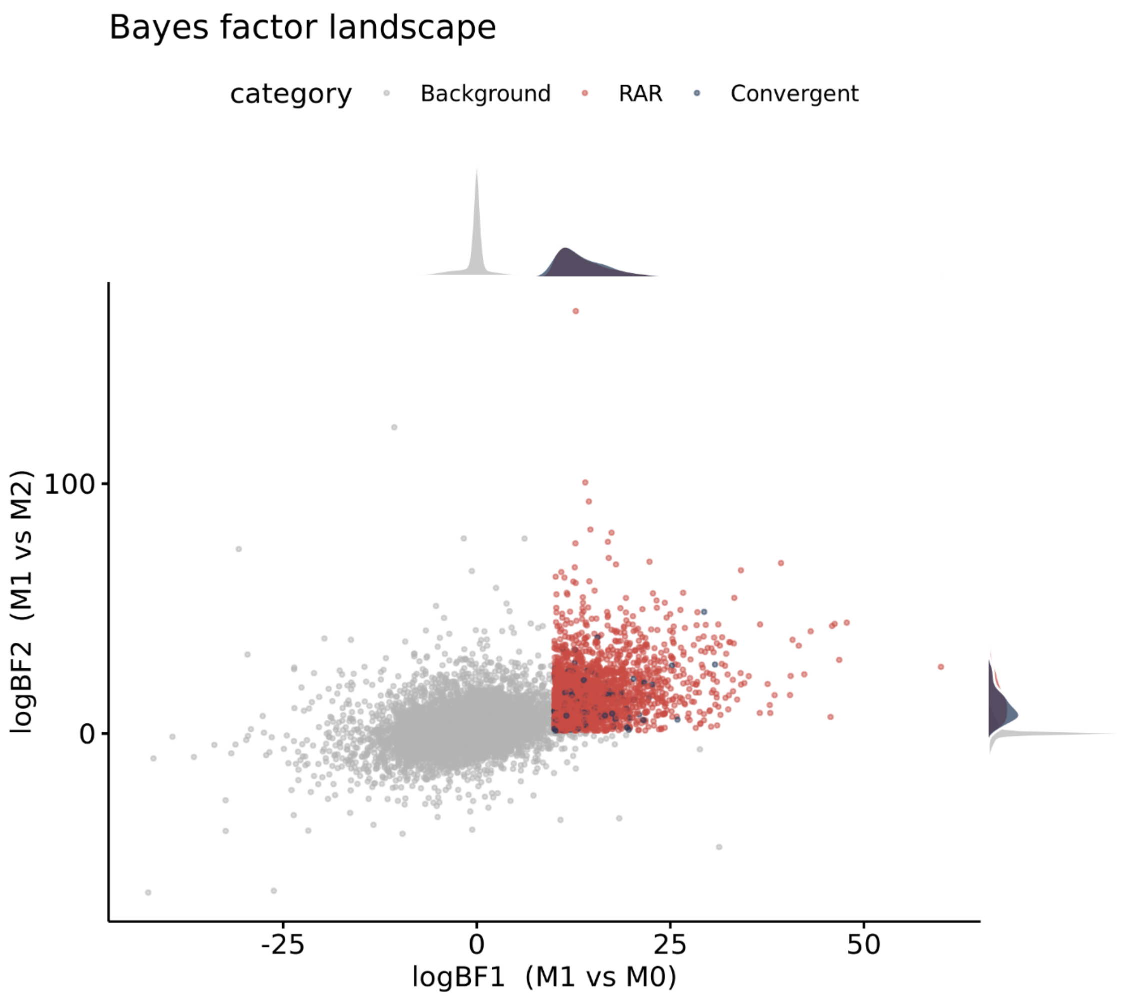


**Fig. S20. Bayes factor landscape of accelerated conserved noncoding elements identified by PhyloAcc-GT.** Scatterplot of conserved noncoding elements (CNEEs) plotted by log Bayes factor support for the target-acceleration model over the conserved null model (logBF1; x-axis) and over the unconstrained acceleration model (logBF2; y-axis). Background CNEEs are shown in gray, lineage-specific endothermic accelerated regulatory regions (EARs) in red, and convergent EARs shared across endothermic lineages in blue.

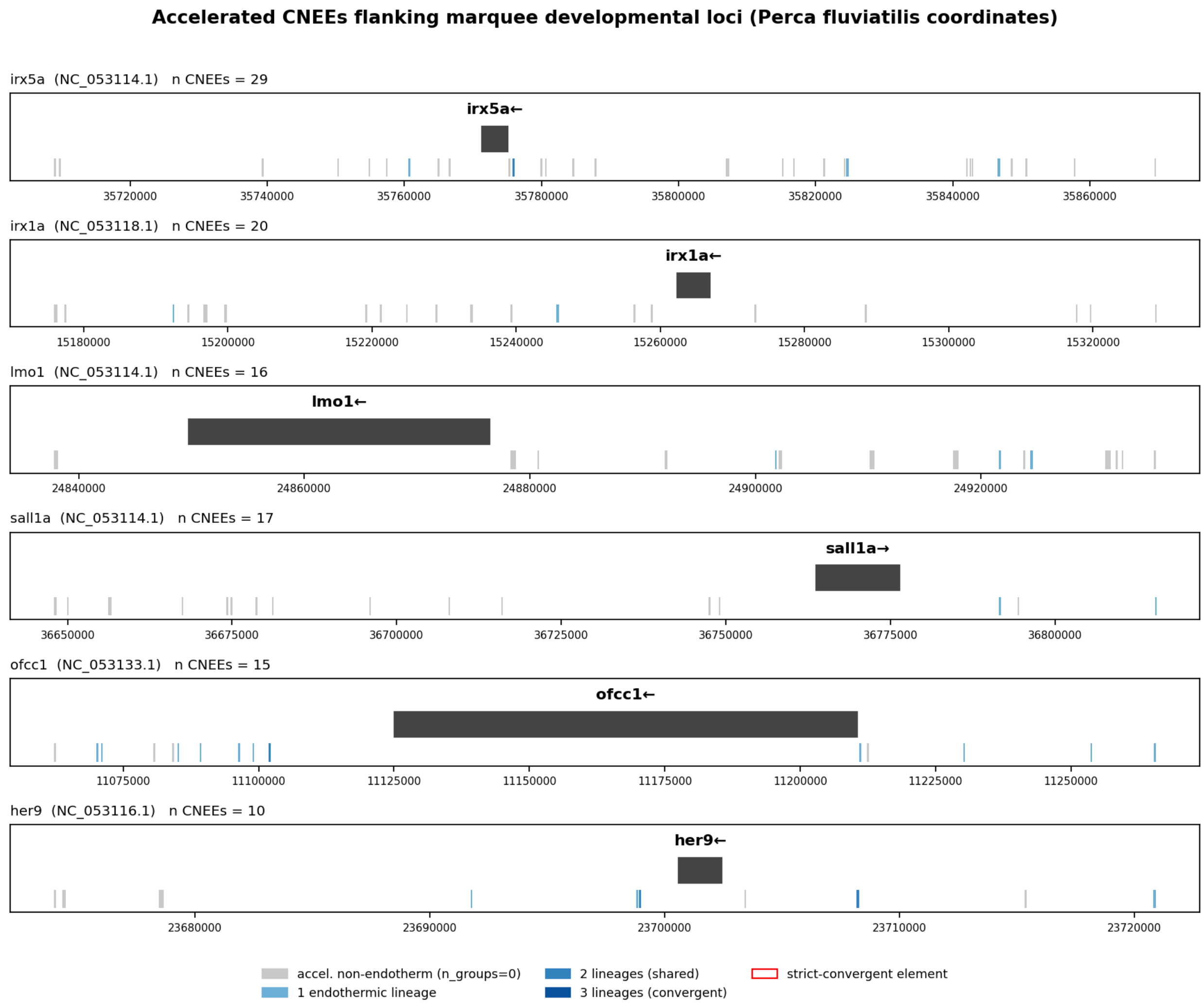


**Fig. S21. In-silico validation of convergent EARs as developmental enhancers.** Accelerated conserved noncoding elements (CNEEs) flanking six marquee developmental loci (*irx5a*, *irx1a*, *lmo1*, *sall1a*, *ofcc1*, *her9*) in *Perca fluviatilis* coordinates; each tick is a CNEE, with grey marking elements accelerated only on non-endothermic lineages and blue shades marking acceleration in one, two, or three endothermic lineages. These loci accumulate many accelerated CNEEs, most on non-endothermic branches, illustrating the regulatory landscape around the convergent developmental regulators. Quantitative results of all validation tests, including the enhancer-mark overlap, are given in Table S21.

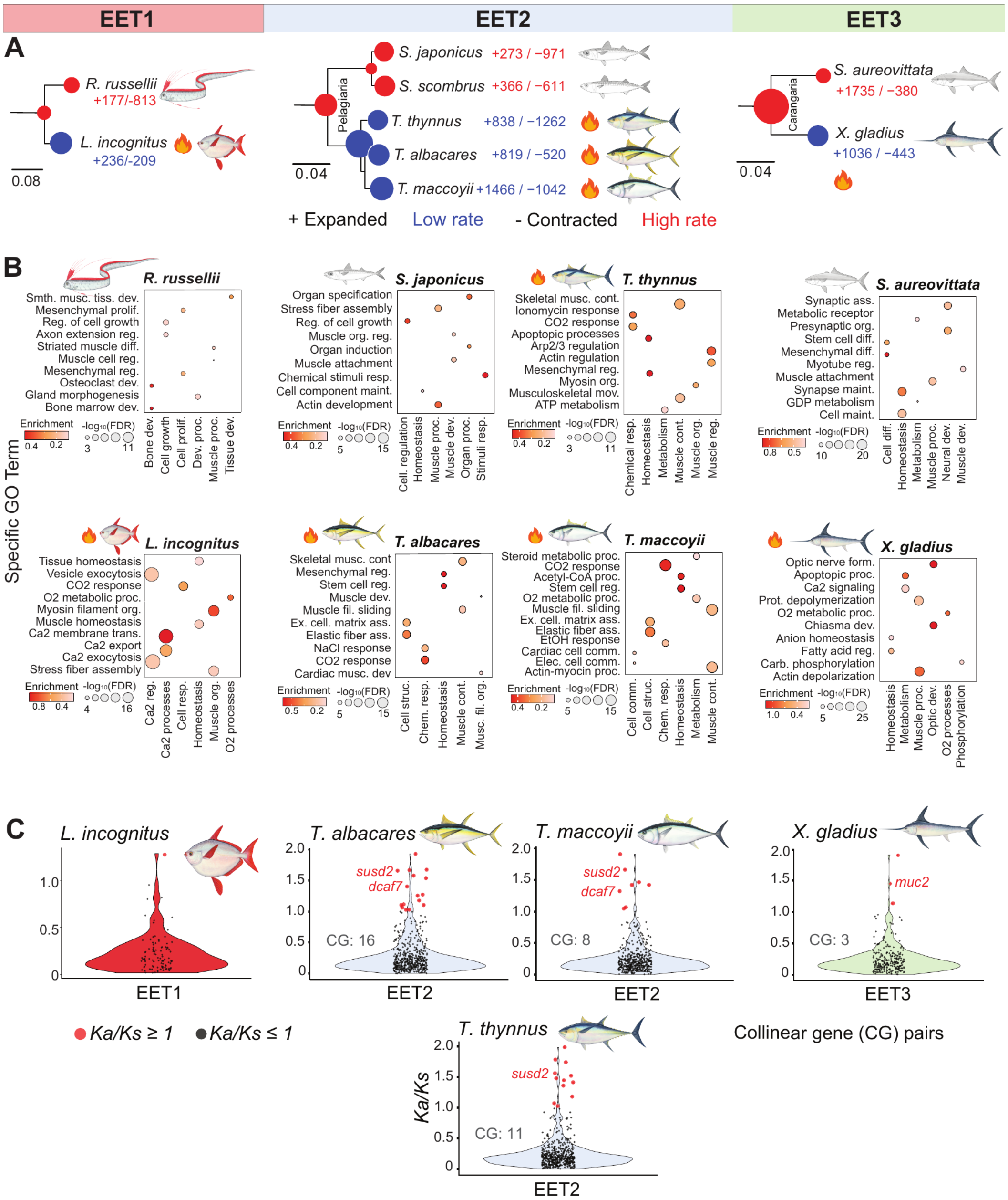

**Fig. S22. Per-lineage gene-family expansion, functional enrichment, and collinear-gene selection (previously Figure 4).** (A) Subtrees for each ectothermy-to-endothermy transition (EET1–EET3). Numbers beside each tip give counts of significantly expanded (+) and contracted (−) gene families along that lineage, from 22,665 tested families; flame symbols mark endothermic species. The ectothermic Carangaria *Seriola dumerili* is omitted because of assembly limitations. Scale bars, substitutions per site. (B) Gene Ontology biological-process enrichment for significantly expanded gene families in each species, grouped by transition. Color encodes the enrichment ratio, and point size encodes statistical support (−log10 FDR); per-species GO terms are on the y-axis and broader functional categories on the x-axis. Enrichment is reported per lineage: each endothermic lineage independently recovered similar broad categories (muscle structure and contraction, calcium homeostasis, oxidative and steroid metabolism, and cellular stress response), but the same categories were also enriched in their ectothermic sisters and the specific expanded families differed among lineages, so these terms reflect parallel functional recruitment rather than convergence across transitions. Full results in Figs. S9–S12 and Tables S6–S11. (C) Distributions of Ka/Ks for collinear orthologous gene pairs (MCScanX) in four endothermic lineages (*T. thynnus*, *T. albacares*, *T. maccoyii*, *X. gladius*). Each point is one collinear gene pair (red, Ka/Ks $\geq 1$; black, Ka/Ks $< 1$). Counts above each violin give the number of candidate pairs per lineage; named candidate loci (*susd2*, *dcaf7*, *muc2*) are labeled.

### Supplementary Tables

**Table S1.** Assembly statistics for the Pacific oarfish (*Regalecus russellii*) genome. Scaffold- and contig-level metrics, gap composition, base composition, masking statistics, and BUSCO completeness.

**Table S2.** List of species used in this study. Order, family, genus, and species of each studied individual alongside NCBI accession numbers and source institutions.

**Table S3.** Mutation statistics for all species in the comparative genomic framework. HAL-based whole-genome alignment metrics per species pair: genome length, transitions and transversions, gap statistics, insertion/deletion events, inversions, duplications, and transpositions.

**Table S4.** Duplication event counts by class across focal endothermic and ectothermic ray-finned fishes (singleton, dispersed, proximal, tandem, and whole-genome/segmental).

**Table S5.** CAFE5 gene-family size evolution across taxa and ancestral nodes. Each row: a gene family (Family ID) and its corresponding taxon or ancestral node, with current copy number, net change, and per-family p-value.

**Table S6.** GO enrichment results for significantly expanded gene families in *Regalecus russellii*. GO term ID, observed and background gene ratios, enrichment metrics, statistical significance, associated gene IDs, and gene counts.

**Table S7.** GO enrichment results for significantly expanded gene families in *Lampris incognitus*.

**Table S8.** GO enrichment results for significantly expanded gene families in *Thunnus thynnus*.

**Table S9.** GO enrichment results for significantly expanded gene families in *Xiphias gladius*.

**Table S10.** GO enrichment results for significantly expanded gene families in *Scomber japonicus*.

**Table S11.** GO enrichment results for significantly expanded gene families in *Seriola aureovittata*.

**Table S12.** MEME codon-level selection results for carnmt1 across endothermic lineages. Per-codon $\alpha$, $\beta_1$, $\beta^+$, their probabilities, LRT, p-value, branches under selection, false-discovery rate (q), and codon classification.

**Table S13.** MEME codon-level selection results for dcaf6 across endothermic lineages.

**Table S14.** Expanded gene families in endothermic ray-finned fishes. Orthogroups significantly expanded across all three endothermic transitions, with predicted-protein annotations, gene names, and curated functional summaries (n = 24 families).

**Table S15.** Collinear gene-pair Ka/Ks ratios in *Lampris incognitus*. Nonsynonymous (Ka), synonymous (Ks), and *Ka/Ks* for collinear pairs.

**Table S16.** Collinear gene-pair *Ka/Ks* ratios in *Thunnus thynnus*.

**Table S17.** Collinear gene-pair *Ka/Ks* ratios in Thunnus albacares.

**Table S18.** Collinear gene-pair *Ka/Ks* ratios in *Thunnus maccoyii*.

**Table S19.** Collinear gene-pair *Ka/Ks* ratios in *Xiphias gladius*.

**Table S20.** Conserved non-exonic elements (CNEEs) under accelerated evolution in endothermic lineages inferred by PhyloAcc-GT. Per-element log-likelihoods under null/target/full models, log Bayes factors (BF1, BF2), and the best-fitting model.

**Table S21. In-silico validation of endothermic accelerated regulatory regions (EARs).** Each test compares the 577 EARs (or the 67 strict-convergent subset) to background conserved noncoding elements (CNEEs). Four location-based tests (expression, positive selection, developmental-gene proximity, enhancer-mark overlap) were corrected together by Benjamini-Hochberg (BH); only enhancer-mark overlap is significant (q = 0.042). The motif-content scan (3b) and the AlphaGenome variant-effect test were evaluated separately and are not part of the four-test correction. P, permutation P unless noted; n.s., not significant. DE, differentially expressed; PMIDs give data sources.

| Test | CNEE set | metric | fold/effect | P | BH q / note |
|---|---|---|---|---|---|
| 1. Expression – DE-gene proximity (PMID 32942994, 34984407) | 577 EAR | nearest-DE membership | 0.86× | 0.95 | n.s. |
| 1b. Expression – DE magnitude | 577 EAR | \|log2FC\| Mann–Whitney | – | 0.47 | n.s. |
| 2. Positive-selection-gene proximity (PMID 40561012) | 577 EAR | broad set (≥1 scenario) | 1.18× | 0.42 | n.s. (small set) |
| 3. Developmental-gene proximity (GO; QuickGO) | 577 EAR | near GO dev/Wnt gene | 1.12× | 0.088 | q = 0.18 |
| 3b. Developmental TF motif content (JASPAR) | 577 EAR | motif-positive, length-matched | 0.79× | 0.96 | n.s. |
| 4. Enhancer-mark overlap (DANIO-CODE; PMID 35789323) | 577 EAR | overlap H3K27ac/H3K4me1 (danRer11) | 1.37× (25.5 vs 18.6%) | **0.010** | **q = 0.042 (significant)** |
| AlphaGenome variant-effect (separate; PMID 41606153) | 67 conv. | endothermic vs. ectothermic-sister pairs, Mann–Whitney | Cliff δ ≈ 0.04 | 0.09–0.11 | n.s. (OOD; see text) |

### Supplementary Datasets

Supplied as separate archive files at the project Figshare repository (doi: pending) Titles and descriptions are provided here for reference.

**Dataset S1.** Estimated phylogenies used in this study. All inferred phylogenetic trees based on chromosome-level genomes, ultraconserved elements (UCEs), and exon markers, provided in Newick format and used for species relationship inference and downstream evolutionary analyses.

**Dataset S2.** Example scripts to generate karyotype files for pairwise synteny visualisation, demonstrating karyotype-file generation from GFF annotations and genome assemblies for use with Circa.

**Dataset S3.** Example scripts to generate synteny .links files for pairwise synteny comparisons.

**Dataset S4.** Input files for MCScanX-based synteny analyses across EET2 and EET3. Collinearity (.collinearity) and GFF3 (.gff) files for all species involved in EET2 and EET3, used as primary input for genome-wide synteny analysis with JCVI/MCScanX.

**Dataset S5.** Input files for microsynteny comparisons of candidate genes (carnmt1, dcaf6). Subdirectories for each gene contain BED files (.bed), gene anchor lists (.anchors), and lifted anchor coordinates (lifted.anchors) for visualising microsynteny around focal genes. Filenames correspond to the species represented.

**Dataset S6.** Input script for PhyloAcc-GT analyses. Primary configuration and command-line scripts used to run PhyloAcc-GT on conserved non-exonic elements (CNEEs).

**Dataset S7.** Python script for PhyloAcc-GT result summarisation and visualisation. Custom Python script used to parse PhyloAcc-GT output and generate summary plots, including Bayes factor distributions and model classifications across accelerated elements; also includes HiSSE-based ancestral state reconstructions used in Fig. S3.
